# Powerful detection of polygenic selection and evidence of environmental adaptation in US beef cattle

**DOI:** 10.1101/2020.03.11.988121

**Authors:** Troy N. Rowan, Harly J. Durbin, Christopher M. Seabury, Robert D. Schnabel, Jared E. Decker

**Author notes:** Corresponding Author: Jared E. Decker.

## Abstract

Selection on complex traits can rapidly drive evolution, especially in stressful environments. This polygenic selection does not leave intense sweep signatures on the genome, rather many loci experience small allele frequency shifts, resulting in large cumulative phenotypic changes. Directional selection and local adaptation are actively changing populations; but, identifying loci underlying polygenic or environmental selection has been difficult. We use genomic data on tens of thousands of cattle from three populations, distributed over time and landscapes, in linear mixed models with novel dependent variables to map signatures of selection on complex traits and local adaptation. We identify 207 genomic loci associated with an animal’s birth date, representing ongoing selection for monogenic and polygenic traits. Additionally, hundreds of additional loci are associated with continuous and discrete environments, providing evidence for local adaptation. These candidate loci highlight the nervous system’s possible role in local adaptation. While advanced technologies have increased the rate of directional selection in cattle, it has likely been at the expense of historically generated local adaptation, which is especially problematic in changing climates. When applied to large, diverse cattle datasets, these selection mapping methods provide an insight into how selection on complex traits continually shapes the genome. Further, by understanding the genomic loci implicated in adaptation, may help us breed more adapted and efficient cattle and begin understanding the basis for mammalian adaptation, especially in changing climates. These selection mapping approaches help clarify selective forces and loci in evolutionary, model, and agricultural contexts.

**Author Summary:** Interest in mapping the impacts of selection and local adaptation on the genome is increasing due to the novel stressors presented by climate change. Until now, approaches have largely focused on mapping “sweeps” on large-effect loci. Highly powered datasets that are both temporally and geographically distributed have not existed. Recently, large numbers of beef cattle have been genotyped across the United States, including influential individuals with cryopreserved semen. This has created multiple powerful datasets distributed over time and landscapes. Here, we map the recent effects of selection and local adaptation in three cattle populations. The results provide insight into the biology of mammalian adaptation and generate useful tools for selecting and breeding better-adapted cattle for a changing environment.

## Introduction

As climate changes, organisms either migrate, rapidly adapt, or perish. The genes and alleles that underlie adaptation have been difficult to identify, except for a handful of large-effect variants that underwent selective sweeps [1]. It is becoming increasingly apparent that for adaptation, hard sweeps are likely to be the exception, rather than the rule [2]. Polygenic selection on complex traits can cause a significant change in the mean phenotype while producing only subtle changes in allele frequencies throughout the genome [3]. Additionally, we expect that polygenic selection is the major selective force both during and after domestication in agricultural species. Many selection mapping methods rely on allele frequency differences between diverged or artificially defined populations (e.g. F_ST_, FLK, XP-CLR) [4–6], making the detection of selection within a largely panmictic population difficult. Others rely on detecting the disruption of normal LD patterns (iHS, EHH, ROH, etc.) [7–9]. In cattle, these methods have successfully identified genomic regions under selection that control Mendelian and simple traits like coat color, the absence of horns, or large-effect genes involved in domestication [10–15]. Further, in many cases these models are unable to derive additional power from massive increases in sample size [16]. Millions of North American *Bos taurus* beef cattle have been exposed to strong artificial and environmental selection for more than 50 years (∼10 generations) [17], making them a powerful model for studying the impacts selection has on genomes over short time periods and across diverse environments.

Though the first cattle single nucleotide polymorphism (SNP) genotyping assay was developed just over a decade ago [18], numerous influential males who have been deceased for 30 to 40 years have been genotyped from cryopreserved semen (Fig S1, Table S1). These bulls add a temporally-stratified, multi-generational component to the datasets of thousands of contemporary animals genotyped from the numerically largest US beef breeds. Furthermore, the large number of animals genotyped from the most recent generations provide remarkable power for detecting small allele frequency changes due to ongoing selection. Under directional selection, alleles will be at significantly different frequencies in more recent generations compared with distant ones (Fig 1c). This creates a statistical association between allele frequencies at a selected locus and an individual’s generation number. With multiple generations sampled and genotyped, we can disentangle small shifts in allele frequency due to directional selection from the stochastic small changes caused by drift (Fig 1a) using Generation Proxy Selection Mapping (GPSM) [17, 19]. The GPSM method searches for allele frequency changes by identifying allelic associations with an individual’s generation (or some proxy), while accounting for confounding population and family structure with a genomic relationship matrix (GRM) (Fig 1d) [20]. When pedigrees are missing, are missing large amounts of data, or have complex, overlapping generations, a proxy for generation can be used such as variety release date or birth date. Cattle producers are selecting on various combinations of growth, maternal, and carcass traits, but the genomic changes that result from this selection are not well-understood. Numerous genome-wide association studies have been undertaken on individual traits, but these say nothing about the underlying genomic changes that populations are experiencing due to selection. The GPSM method identifies the allele frequency changes underlying selection in a trait-agnostic manner, allowing us to observe the impacts of selection decisions in real time and understand how strong selection alters the genome over very short timescales.

**Fig 1.**
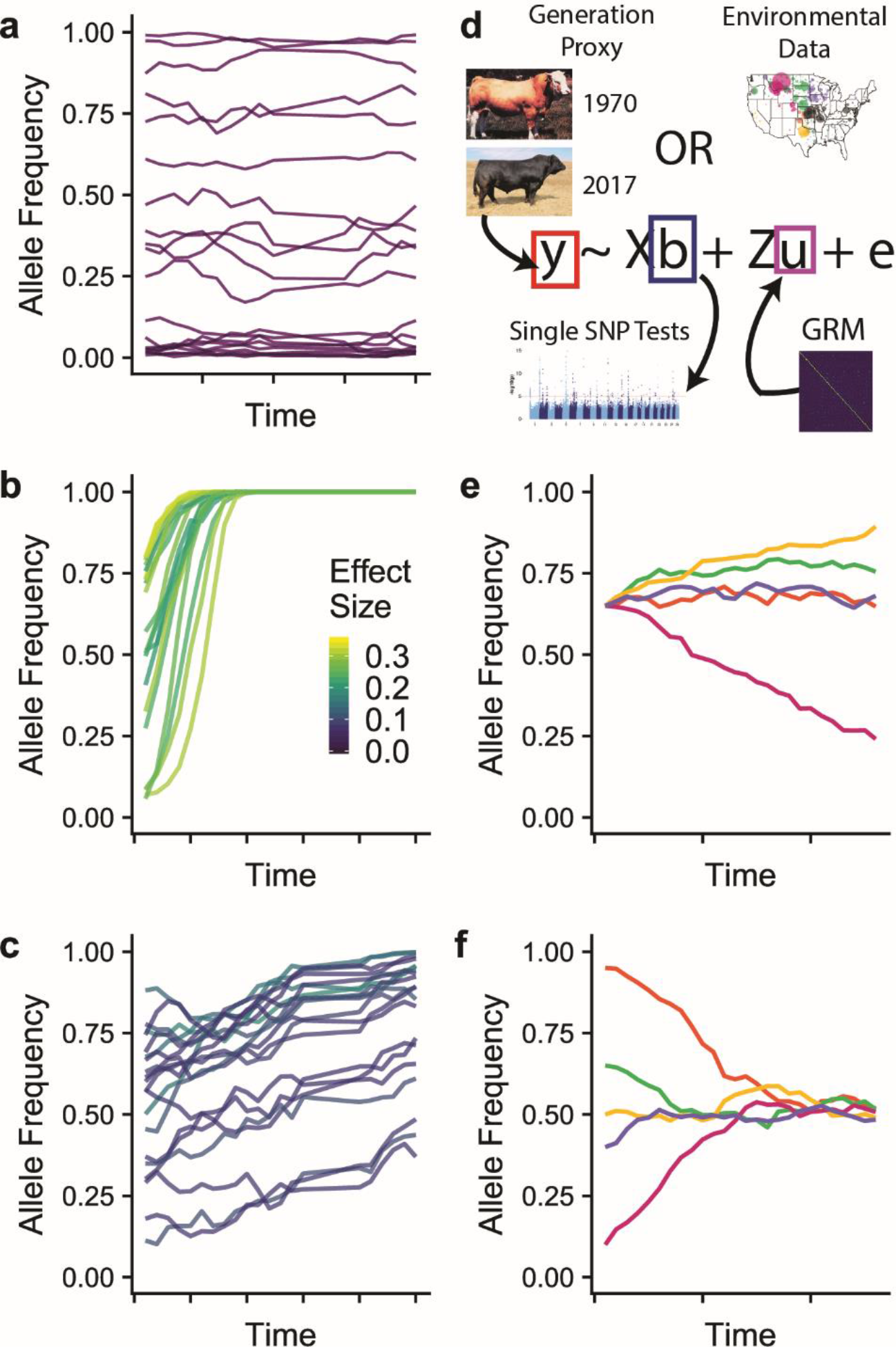
Simulated allele frequency trajectories and model overview. (a-c) Allele frequency trajectories for 20 SNPs colored by relative effect sizes from stochastic selection simulations. (a) Effect size = 0, representing stochastic changes in allele frequency due to genetic drift. (b) Large-effect alleles rapidly becoming fixed in the population representing selective sweeps. (c) Moderate-to-small effect size SNPs changing in frequency slowly over time, representing polygenic selection. (d) An overview of the linear mixed model approach used for Generation Proxy Selection Mapping and environmental GWAS. (e-f) A single SNP under ecoregion-specific selection. Different colors represent the trajectory of a given SNP in one of five different ecoregions. Ecoregion-specific selection can lead to allele frequencies that (e) diverge from or (f) converge to the population mean.

**Table 1.**
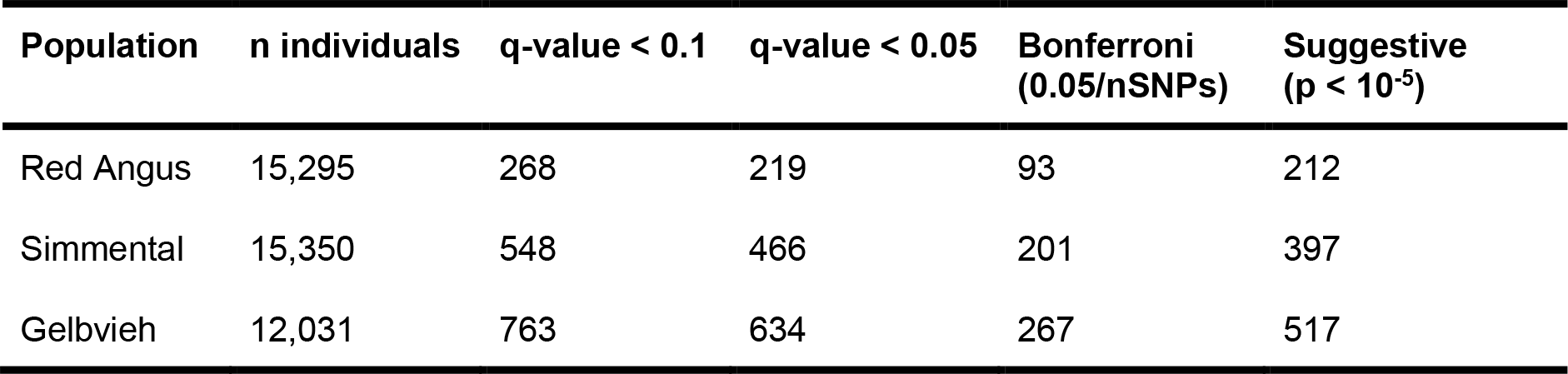
Number of significant birth date-associated SNPs in each population at various significance thresholds.

| Population | n individuals | q-value < 0.1 | q-value < 0.05 | Bonferroni<br>(0.05/nSNPs) | Suggestive<br>($p < 10^{-5}$) |
| --- | --- | --- | --- | --- | --- |
| Red Angus | 15,295 | 268 | 219 | 93 | 212 |
| Simmental | 15,350 | 548 | 466 | 201 | 397 |
| Gelbvieh | 12,031 | 763 | 634 | 267 | 517 |

From domestication to the present, humans have used phenotypic selection to change cattle. Since the 1980s and since the 2010s, genetic and genomic predictions, respectively, have been available to U.S. beef producers. However, even with these advanced tools, most beef producers still rely, at least partially, on phenotypic selection [21]. Prior to the 1980s, cattle were selected via phenotypic selection and there was very little movement of animals (i.e. gene flow) between regions. This strong, artificial phenotypic selection allows unintended selection on naturally-occurring abiotic and biotic stressor traits, akin to natural selection. Further, phenotypic selection could act on loci with genotype-by-environment effects (which BLUP breeding value-based selection would not), thus creating local adaptation. As an example, phenotypic selection could select for animals with better innate immune systems as they will grow faster and look more vigorous.

Though domesticated, beef cattle are exposed to a broad spectrum of unique environments and local selection pressures, as compared to other more intensely managed livestock populations. This suggests that local adaptation and genotype-by-environment interactions play important roles in the expression of complex traits. Local adaptation and genotype-by-environment (GxE) interactions are known to exist in closely related cattle populations [22, 23]. Previous work has identified the presence of extensive GxE in beef cattle populations [24–28], but limited work exploring the genomic basis of local adaptation has occurred [29]. Further, we anticipate that the increased use of artificial insemination, responsible for dramatic increases in production efficiency, may be eroding environmentally adaptive allele frequency differences in populations. Understanding genetic interactions with the environment, and their presence in cattle populations will become increasingly important in the face of changing climates.

To identify genomic regions potentially contributing to local adaptation, we used continuous environmental variables as quantitative phenotypes or discrete ecoregions as case-control phenotypes in a linear mixed model framework. We refer to these approaches as “environmental genome-wide association studies”, or envGWAS. Using a genomic relationship matrix in a LMM allows us to control the high levels of relatedness between spatially close individuals, and more confidently identify real environmentally-associated alleles. This method builds on the theory of the Bayenv approach from Coop et al. (2010) [30, 31] that uses allele frequency correlations along environmental gradients to identify potential local adaptation.

Herein, we use two methods (Fig 1), the first for detecting complex polygenic selection (Generation Proxy Selection Mapping, GPSM), and the second for identifying local adaptation (environmental Genome-Wide Association Studies, envGWAS). When applied to three US beef cattle populations, each with ∼15,000 genotyped individuals, we identified numerous genomic regions harboring directional or environmentally associated mutations. Further, using a meta-analysis approach, we identified loci responding to region-specific selection (Fig 1e,f), largely due to the erosion of local adaptation caused by gene flow among ecoregions from the use of artificial insemination sires. This study is the first step in assisting beef cattle producers to identify locally adapted individuals, which will reduce the industry’s environmental footprint by increasing efficiency and resilience to stressors. Further, this repurposing of commercially-generated genomic data provides us unprecedented power to gain insight into the biology of polygenic selection and adaptation in mammalian species.

## Results

### Simulations

To identify the robustness of GPSM to distinguish selection from drift, we performed two major sets of simulations, stochastic and gene drop. First, we performed a set of stochastic simulations to demonstrate how selection in the context of different effective population sizes, selection intensities, generational sampling, and genomic architectures produce GPSM signal.

Stochastic simulations under multiple selection intensities, time periods, and trait architectures showed consistently that GPSM is able to map polygenic selection (**Table S2**). Across all simulated scenarios and architectures, we identified an average of 38.5 selected loci (min 5.2, max 64.1) that reached genome-wide significance (q < 0.1) in GPSM. Depending on the genomic architecture of the simulated trait under selection, this represented between 0.57% and 32.54% of possible true positives (median = 9.94%). In most cases, we observed that significant hits were not the largest effect simulated QTL, but the loci that underwent the greatest allele frequency shifts over the course of genotype sampling. Usually, the largest effect QTL were fixed in the population during burn-in simulations, making their detection by GPSM impossible. Simulations suggested that GPSM effectively distinguished allele frequency changes due to selection from those associated with drift. Across all 36 scenarios of random selection, with drift acting as the only force by which alleles could change in frequency, we detected an average of 0.15 GPSM false positive SNPs (q-value < 0.1) per replicate. These rare false positives were not driven by changes in any single component of the simulations. The average number of false positive SNPs detected per replicate ranged from 0.0 to 0.45 across all scenarios, accounting for, at most, 1 false positive GPSM SNP per 100,000 tests.

**Table 2.** Variation in birth date explained by three classes of SNPs. The PVE estimates (standard error in parentheses) from a genomic restricted maximum likelihood (GREML) variance component analysis of birth date using three GRMs created from: 1) genome-wide significant SNPs (q < 0.1), 2) an equivalent number of the next most significant SNPs outside of genome-wide significant associated regions, and 3) an equivalent number of randomly sampled SNPs from genomic regions that did not harbor genome-wide significant associations.

| Population | Genome-wide significant SNPs* | Suggestive significant SNPs* | Other SNPs* | Total |
| --- | --- | --- | --- | --- |
| Red Angus | 0.170 (0.021) | 0.160 (0.016) | 0.030 (0.004) | 0.360 (0.020) |
| Simmental | 0.223 (0.017) | 0.199 (0.014) | 0.046 (0.004) | 0.468 (0.015) |
| Gelbvieh | 0.232 (0.018) | 0.175 (0.012) | 0.021 (0.003) | 0.428 (0.016) |

\*Contained 268, 548, and 763 SNPs for Red Angus, Simmental, and Gelbvieh, respectively

Genotype sampling also significantly impacted the number of true and false positives detected by GPSM across scenarios. When genotypes were evenly sampled across generations, GPSM detected, on average, 18.05 more true positives (paired t-test p < 2 x 10^-16^) and 0.90 less false positives (paired t-test p < 2 x 10^-16^) compared with heavier sampling of recently born individuals. That said, across all scenarios, the uneven sampling scheme that more closely resembled our real datasets still detected over 20 selected loci on average across all simulations (min 1.9, max 36.4) The proportion variation explained (PVE, **Table S2**) of birth date across simulation scenarios reflects the number of generations and the number of crosses in the simulations across both selection and random scenarios (p < 2e-16). For random scenarios, the number of segregating QTL (p = 9.94e-06) was also associated with PVE. Importantly, for selection scenarios, the proportion of true positives (p = 2.87e-07) and QTL distribution (p = 2.91e-13) were also significantly associated with the PVE.

Gene dropping simulations using the Red Angus pedigree generated an average of 0.4 (sd = 0.52) significant GPSM loci (q < 0.1) per 200K markers tested, equating to 1-2 in our real genotype data (Table S1). Thus, pedigree structure is not responsible for the significant SNPs that we detected in real datasets.

### Detecting ongoing polygenic selection with Generation Proxy Selection Mapping (GPSM)

We used continuous birth date and high-density SNP genotypes for large samples of animals from three large US beef cattle populations; Red Angus (RAN; n =15,295), Simmental (SIM; n =15,350), and Gelbvieh (GEL; n =12,031) to map loci responding to polygenic selection (Table S3, Fig S1). The LMM estimated that the proportion of variance in individuals’ birth dates explained by the additive genetic effects of SNPs was large [Proportion of Variance Explained (PVE) = 0.520, 0.588, and 0.459 in RAN, SIM, and GEL, respectively], indicating that the demographic histories of the populations and sampling strategy across the breeds were similar. We removed the link between generation proxy and genotype by randomly permuting the animals’ birth date and on reanalysis of the permuted data we observed PVE to decrease to zero (Table S4).

The GPSM analyses for these three populations identified 268, 548, and 763 statistically significant SNPs (q-value < 0.1), representing at least 52, 85, and 92 genomic loci associated with birth date in RAN, SIM, and GEL, respectively (Fig 2a-f, Table 1, **Table S5**). Additionally, we found that despite birth date being a significantly skewed dependent variable (RAN skewness = 4.48, SIM skewness = 2.66, GEL skewness = 3.64), the GPSM p-values were well calibrated (Fig 2g-i). Despite the tendency for genome-wide association studies (GWAS) to be biased in its detection of moderate frequency variants [32], we identify significant associations across the minor allele frequency range in our GPSM simulations and analyses (Fig 2j-l). This suggests GPSM can differentiate drift from selection across the allele frequency spectrum. Rapid shifts in allele frequency create highly significant GPSM signals. For example, *rs1762920* on chromosome 28 has undergone large, recent changes in allele frequency in all three populations (Fig 2g), which in turn creates highly significant q-values (2.810⨯10^-27^, 2.323⨯10^-150^, 2.787⨯10^-265^ in RAN, SIM, and GEL, respectively). The allele frequency changes observed for this locus are extremely large compared to other significant regions, most of which have only small to moderate changes in allele frequency over the last ∼10 generations. When we regressed allele frequency (0, 0.5, or 1.0 representing *AA*, *AB*, and *BB* genotypes per individual) on birth date, the average allele frequency changes per generation (ΔAF) for significant GPSM associations were 0.017, 0.024, and 0.022 for RAN, SIM, and GEL, respectively (Table S6). In the analyses of each dataset, GPSM identified significant SNPs with ΔAF < 1.1⨯10^-4^. The generally small allele frequency changes detected by GPSM are consistent with the magnitude of allele frequency changes expected for selection on traits with polygenic architectures [3].

**Fig 2.**
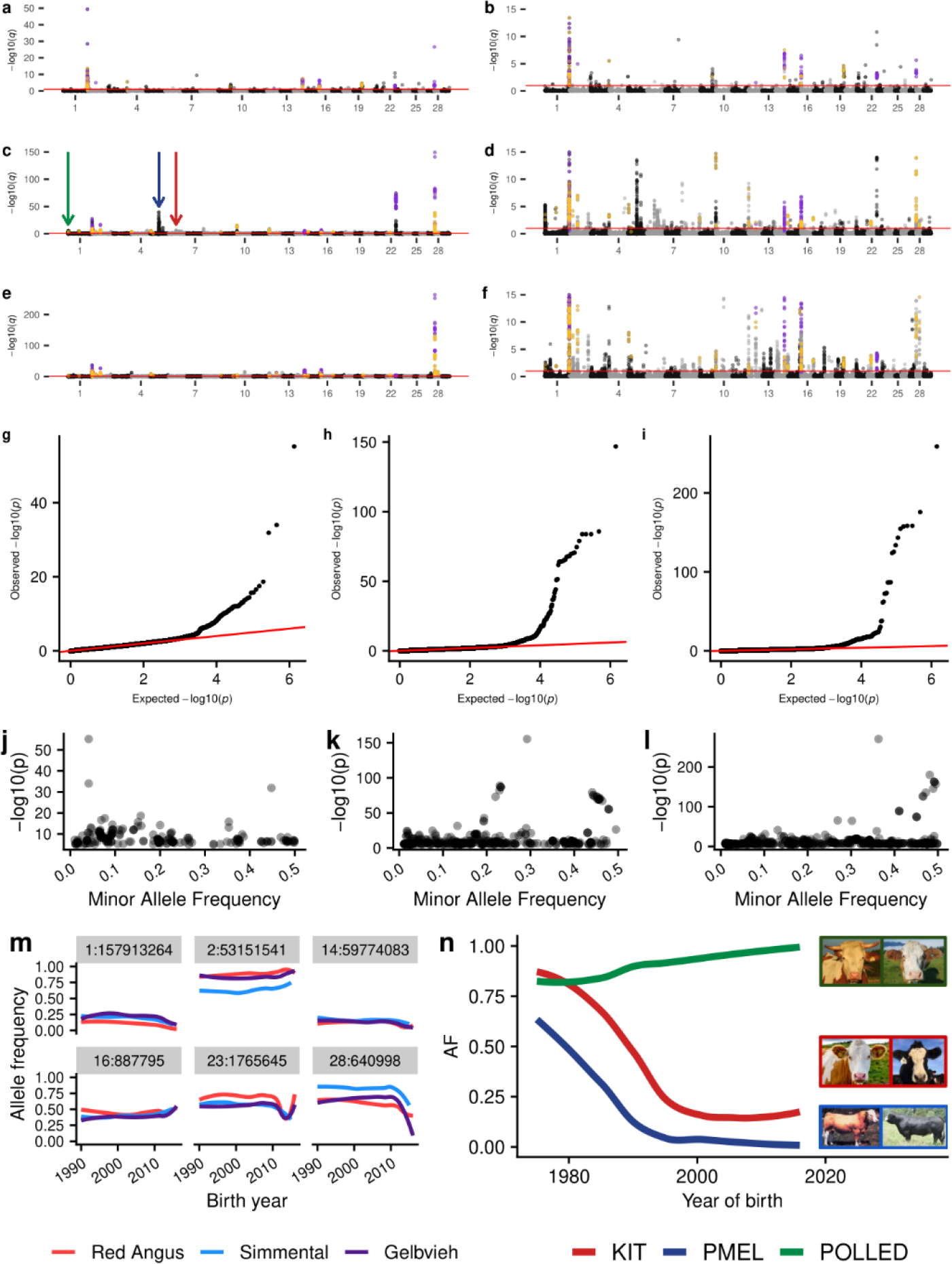
Generation Proxy Selection Mapping identifies signals of polygenic selection in three major U.S. cattle populations. Full and truncated (-log_10_(*q*)< 15) Manhattan plots for GPSM analysis of Red Angus (a & b), Simmental (c & d), and Gelbvieh (e & f). Purple points indicate SNPs significant in all three population-specific GPSM analyses and orange points indicate SNPs significant in two. Red lines in a-f indicate a significance cutoff of q < 0.1. Quantile-quantile plots of -log_10_(*p*) values in GPSM analysis of (g) Red Angus, (h) Simmental, and (i) Gelbvieh populations. Minor allele frequency plotted versus -log10(*p*) values for significant SNPs in (j) Red Angus, (k) Simmental, and (l) Gelbvieh populations. (m) Smoothed allele frequency histories for the six most significant loci identified as being under selection in all three datasets. (n) Allele frequency histories for three known Mendelian loci that control differences in visual appearance between introduced European and modern US Simmental cattle. Arrows of the same color are used to distinguish the genomic locations of these loci in (c).

We found that significant GPSM loci go largely undetected when using other selection mapping methods. In these datasets, the genomic regions identified by both TreeSelect and allele frequency trajectory (wfABC) methods for detecting selection were almost entirely different from those identified by GPSM. Further, the estimated selection coefficients (*s*) from wfABC [33] were lowly correlated with GPSM effect size estimates using both the full (*r* = 0.002) and relationship-pruned (*r* = 0.003) datasets. The slight increase in correlation is likely due to a violation of wfABC’s assumption of random mating, because the correlation between estimated *s* between the full and relationship-pruned datasets was only 0.432. When we restricted our comparison to significant GPSM SNPs, the correlation between wfABC-estimated *s* and GPSM betas was higher (*r* = 0.0675), but still quite low. Interestingly the correlation between estimated selection coefficients and significant GPSM p-values was higher than with the GPSM effect size estimate (*r* = 0.126).

TreeSelect and GPSM test statistics were also lowly correlated. A three population TreeSelect analysis that treated each breed as a distinct population to detect between-breed selection showed low correlations between test statistics (*r* = 0.009, 0.002, 0.002 in Red Angus, Simmental, and Gelbvieh, respectively). The between-breed allele frequency differences detected by TreeSelect did not overlap at all with GPSM signatures. When used to detect selection within a breed using discrete groupings based on animal birth date, test statistic correlations were also low (*r* = 0.007, 0.018, and 0.005 for youngest, middle, and oldest groups of animals in Red Angus). Despite generally low correlations between test statistics, TreeSelect with arbitrary population groupings detected some of the same loci that were identified by GPSM. In Red Angus, the within-breed TreeSelect analysis identified selection at six loci that were also significant in GPSM, including the two most significant loci on chromosome 23 near *KHDRBS2* and on chromosome 28 near *RHOU* (Fig S2).

We performed a genomic restricted maximum likelihood (REML) analysis to identify how much of the variation in birth date was explained by various classes of GPSM SNPs. We built three GRMs using different SNP sets: One set with GPSM genome-wide significant SNPs (q < 0.1), the second with an equivalent number of the next most suggestive GPSM SNPs outside of significant loci (> 1 Mb from a q < 0.1 significant SNP), and the third with an equivalent number of random, moderate minor allele frequency (MAF > 0.15) SNPs not in the first two variant classes, intended to represent loci randomly drifting in the population. For each population, we observed that nearly all of the variation in birth date was explained by the significant and suggestive GRMs. While genome-wide significant loci explain the majority of genetic variance associated with birth date, an equivalent number of suggestive, but not significant SNPs resulted in only slightly smaller PVEs (Table 2). We suspect that these SNPs are undergoing directional allele frequency changes too small to detect at genome-wide significance, even in this highly-powered dataset. Since GPSM continues to gain power with additional samples, we suspect that future sample size increases will detect more of these signatures of polygenic selection at a genome-wide significance level. Regardless of the number of SNPs used in the drift GRM, the variance associated with drift was consistently minimal (Table 2).

As proof-of-concept, GPSM identified known targets of selection. In Simmental, we identified significant associations at three known Mendelian loci that explain the major differences in appearance between early imported European Simmental and modern US Simmental (Fig 2h). These loci: *POLLED* (absence of horns [34]), *ERBB3/PMEL* (European Simmental cream color [35]), and *KIT* (piebald coat coloration [36]) have not appreciably changed in allele frequency since 1995, making their GPSM signature significant, but less so than other loci actively changing in frequency.

In addition to these three known Mendelian loci, we detected numerous novel targets of selection within and across populations. While the majority of the genomic regions detected as being under selection were population-specific (79.8%, 79.8%, and 77.2% of the significant regions in RAN, SIM, and GEL, respectively), we identified seven loci that are under selection in all three populations, and fifteen more under selection in two (Table S7). While GPSM is able to detect Mendelian selection, the overwhelming majority of signatures identified represent selection on complex, quantitative traits. Of the regions identified in multiple populations, many possessed positional candidate genes with production-related functions in cattle (*DACH1*-Growth [37, 38], *LRP12*-Growth [39], *MYBPH*-Muscle Growth [40], *RHOU*-Carcass Weight [41], *BIRC5*-Feed Intake [42]). However, GPSM did not identify any of the well-established large-effect growth loci (e.g., chromosome 14 locus containing *PLAG1*, chromosome 6 locus containing *LCORL*). Growth phenotypes (e.g., birth, weaning, and yearling weights) are known to be under strong selection in all three populations [43], but antagonistic pleiotropic effects such as increased calving difficulty prevent directional selection from changing frequencies at these large-effect loci. Many of the selection signatures identified in at least two of the populations have no known functions or phenotype associations in cattle, highlighting the ability of GPSM to identify novel, important loci under polygenic selection, agnostic of phenotype.

Biological processes and pathways enriched in genes located proximal to GPSM SNP associations point to selection on drivers of production efficiency and on population-specific characteristics (**Table S8**). In each population, we identified numerous biological processes involved in cell cycle control, which are directly involved in determining muscle growth rate [44], as being under selection. In Red Angus and Gelbvieh we identified multiple cancer pathways as being under selection. This likely represents further evidence of selection on cell cycle regulation and growth rather than on any cancer related phenotypes [45]. Red Angus cattle are known to be highly fertile with exceptional maternal characteristics [46]. We identified the “ovarian steroidogenesis” pathway as being under selection, a known contributor to cow fertility [47]. We also identify numerous other processes involved in the production and metabolism of hormones. Hormone metabolism is a central regulator of growth in cattle [48], but could also represent selection for increased female fertility in Red Angus. Further, Tissue Set Enrichment Analyses (TSEA) of Red Angus GPSM candidate genes showed suggestive expression differences (p < 0.1) in multiple human reproductive tissues (**Tables S9-S10**). Enrichments in these tissues did not exist in TSEA of Simmental or Gelbvieh GPSM gene sets, suggesting explicit within-population selection on fertility. Gelbvieh cattle are known for their rapid growth rate and carcass yield. Selection on these phenotypes likely drives the identification of the six biological processes identified which relate to muscle development and function in the Gelbvieh GPSM gene set. Consequently, this gene set is significantly enriched for expression in human skeletal muscle (**Tables S9-S10**), an enrichment unique to Gelbvieh. A complete list of genomic regions under population-specific selection and their associated candidate genes is in **Table S5**.

### Detecting environmental associations using envGWAS

Using an equivalent form of model to GPSM, but with continuous environmental variables (30 year normals for temperature, precipitation, and elevation) or statistically-derived discrete ecoregions as the dependent variable (rather than birth date in GPSM) allows us to identify environmentally-associated loci that have been subjected to artificial and, perhaps in this context more importantly, natural selection [49]. We refer to this method as environmental GWAS (envGWAS). envGWAS extends the theory of the Bayenv approach of Coop et al. (2010) which searches for allele frequency correlations along environmental gradients to identify potentially adaptive loci [30]. Similar approaches have been applied to plant datasets [50, 51], but we extend envGWAS to panmictic, biobank-sized mammalian populations. Unlike many genome-environment association analyses which only used linear models [51, 52], our large dataset and the use of multivariate models provides power to identify association while importantly controlling for geographic dependence between samples using a genomic relationship matrix (Fig S3, Fig S7). We used *K*-means clustering with 30-year normal values for temperature, precipitation, and elevation to partition the United States into 9 discrete ecoregions (Fig 3a). These ecoregions are largely consistent with those represented in previously-published maps from the environmetrics and atmospheric science literature [53], and reflect well-known differences in cattle production environments. The resulting ecoregions capture not only combinations of climate and environmental variables, but associated differences in forage type, local pathogens, and ecoregion-wide management differences to which animals are exposed. Thus, using these ecoregions as case-control phenotypes in envGWAS allowed us to detect more complex environmental associations. The three studied populations are not universally present in all ecoregions (Fig 3b, Fig S4b & S5b, Table S11). Since the development of these US populations in the late 1960s and early 1970s, registered seedstock animals from these populations have a small footprint in desert regions with extreme temperatures and low rainfall.

**Fig 3.**
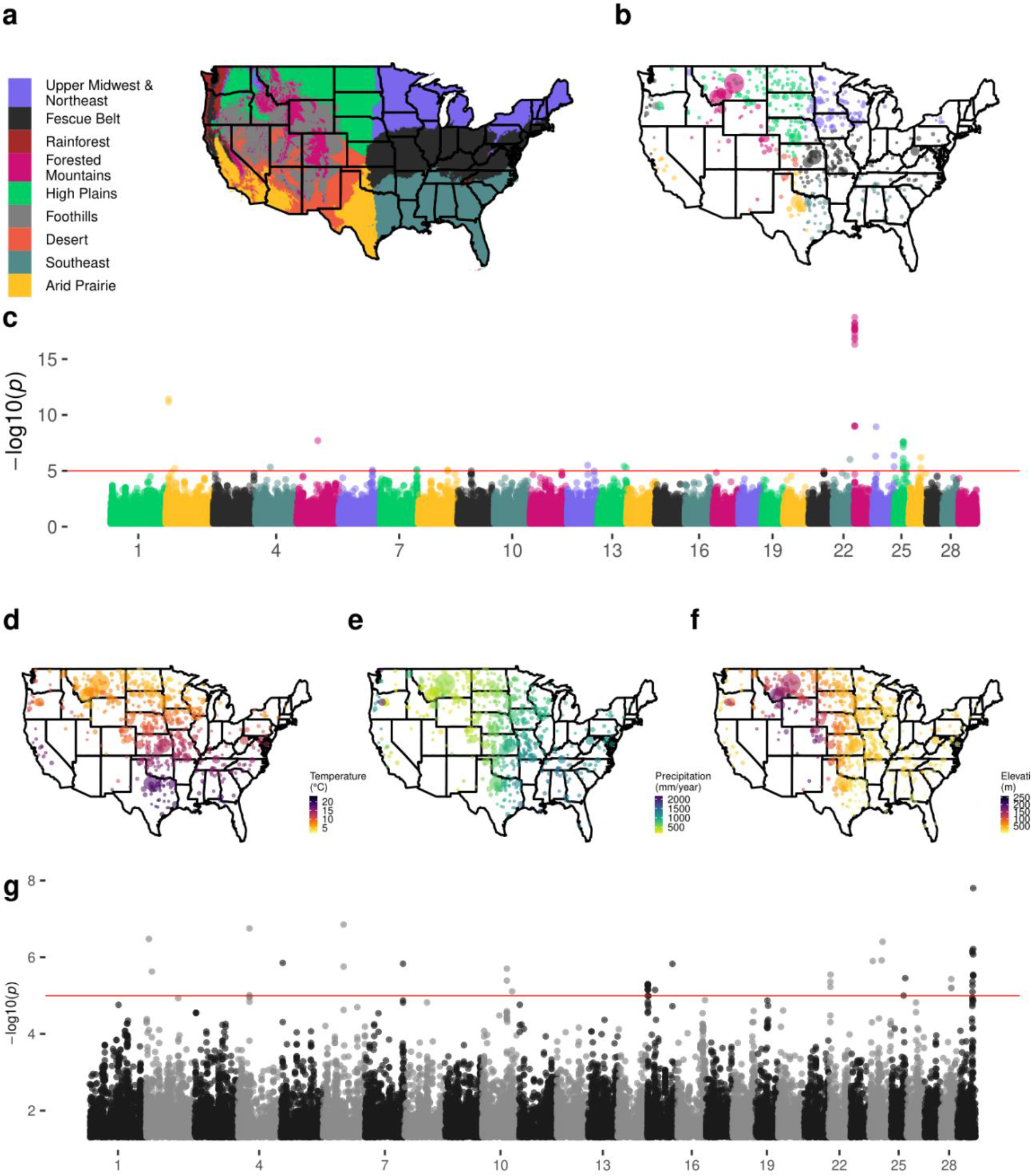
Manhattan plots for discrete and continuous envGWAS in Red Angus cattle. (a) Nine continental US ecoregions defined by *K*-means clustering of 30-year normal temperatures, precipitations, and elevations. (b) Locations of sampled Red Angus animals colored by breeder’s ecoregion and sized by the number of animals at that location. (c) Multivariate discrete envGWAS (case-control for six regions with > 600 animals). Locations of sampled Red Angus animals colored by (d) 30-year normal temperature, (e) 30-year normal precipitation, and (f) elevation. (g) Multivariate continuous envGWAS with temperature, precipitation, and elevation as dependent variables. For all Manhattan plots the red line indicates the empirically-derived p-value significance threshold from permutation analysis (p < 1⨯10^-5^).

Although environmental variables and ecoregions are not inherited, the estimated PVE measures the extent to which genome-wide genotypes change in frequency across the environments in which the animals were born and lived. The proportion of variance explained by SNPs ranged from 0.586 to 0.691 for temperature, 0.526 to 0.677 for precipitation, and 0.585 to 0.644 for elevation (Table S12). In Red Angus, PVE for ecoregion membership ranged from 0.463 for the Arid Prairie to 0.673 for the Fescue Belt (Table S13). We observe similar environmental PVE in both Simmental and Gelbvieh datasets. These measures suggest that genetic associations exist along both continuous environmental gradients and within discrete ecoregions. Permutation tests that shuffled environmental dependent variables, removing the relationship between the environment and the animal’s genotype, resulted in all PVEs being reduced to ∼ 0, strongly suggesting that the detected associations between genotype and environment were not spurious. An additional permutation test that permuted animals’ zip codes, such that all animals from a given zip code were assigned the same “new’’ zip code from a potentially different ecoregion provided similar results, indicating that bias due to sampling at certain zip codes was not producing envGWAS signals. From 10 rounds of permutation, there were no SNP associations with p-values < 1⨯10^-5^. Consequently, we used this empirically-derived p-value threshold to determine SNP significance in all of the envGWAS analyses. Gene drop simulations suggested that a small portion of the identified associations are likely due to pedigree structure or founder effects (average of 1.36 false positive envGWAS loci per 200,000 tests). However, in this data, the pedigree structure reflects selection decisions of farmers and ranchers that are not beyond the influence of performance differences relative to environmental differences.

### Discrete ecoregion envGWAS

In Red Angus, we identified 54 variants defining 18 genomic loci significantly associated with membership of an ecoregion in the discrete multivariate envGWAS analysis (Fig 3c). Despite locus-specific signal, principal component analysis (PCA) does not suggest that ecoregion-driven population structure exists in any of the populations (Fig S6). Of these loci, only two overlapped with loci identified in the continuous envGWAS analyses, suggesting that using alternative definitions of environment in envGWAS may detect different sources of adaptation. Of the 18 significant loci, 17 were within or near (< 100 kb) positional candidate genes (Tables S14-**S15**), many of which have potentially adaptive functions. For example, envGWAS identified SNPs immediately (22.13 kb) upstream of *CUX1* (Cut Like Homeobox 1) gene on chromosome 25. *CUX1* controls hair coat phenotypes in mice [54], and alleles within *CUX1* can be used to differentiate between breeds of goats raised for meat versus those raised for fiber [55]. The role of *CUX1* in hair coat phenotypes makes it a strong adaptive candidate in environments where animals are exposed to heat, cold, or toxic ergot alkaloids from fescue stress [56].

In Simmental, we identified 11 loci tagged by 39 variants significantly associated with membership of an ecoregion in the multivariate envGWAS analysis (Fig S4). In Gelbvieh, 66 variants identified 33 local adaptation loci (Fig S5). In the analyses of all three datasets, we identified a common local adaptation signature on chromosome 23 (peak SNP *rs1023574*). Multivariate analyses in all three populations identified alleles at this SNP to be significantly associated with one or more ecoregions (q = 1.24 x 10^-13^, 3.15 x 10^-12^, 4.82 x 10^-5^ in RAN, SIM, and GEL, respectively). In all three datasets, we identified *rs1023574* as a univariate envGWAS association with membership of the Forested Mountains ecoregion. However, the most significant univariate association in Red Angus was with the Arid Prairie region which was excluded from both the Simmental and Gelbvieh analyses due to low within-region sample size. In the multivariate analysis for Red Angus, the associated locus spanned 18 SNPs from (1,708,914 to 1,780,836 bp) and contained the pseudogene *LOC782044*. The nearest annotated gene, *KHDRBS2* (KH RNA Binding Domain Containing, Signal Transduction Associated 2) has previously been identified by other adaptation studies in cattle, sheep, and pigs [57–59]. This variant was not significantly associated with any continuous environmental variable in Red Angus. However, *rs1023574* was significantly associated with temperature, elevation, and humidity variables in Simmental. The *KHDRBS2* locus was preferentially introgressed between *Bos taurus* and domestic yak [60]. Further, this locus shows an abnormal allele frequency trajectory (Fig 4c), indicating that it may be a target of balancing selection.

**Fig 4.**
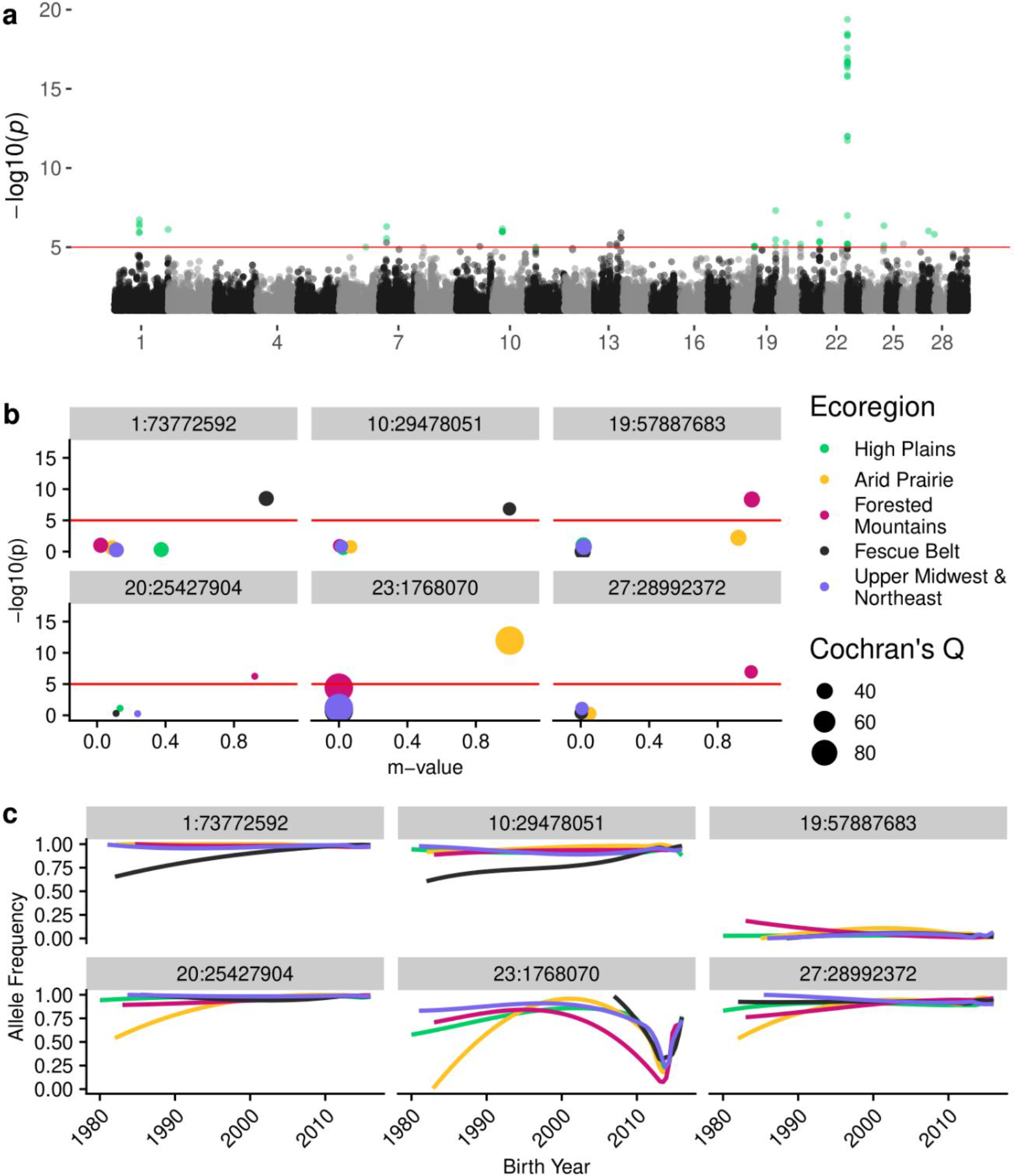
Meta-analysis of within-ecoregion GPSM for Red Angus cattle. (a) Manhattan plot of per-variant Cochran’s Q p-values. Points colored green had significant Cochran’s Q (p < 1⨯10^-5^) and were significant in at least one within-region GPSM analysis (p < 1⨯10^-5^). (b) Ecoregion effect plots for lead SNPs from six loci from (a). Points are colored by ecoregion and are sized based on Cochran’s Q value. (c) Ecoregion-specific allele frequency histories for SNPs from (b), colored by ecoregion.

### Continuous environmental variable envGWAS

Using continuous temperature, precipitation, and elevation data as quantitative dependent variables in a multivariate envGWAS analysis of Red Angus animals, we identified 46 significantly associated SNPs (Fig 3g). These SNPs tag 17 loci, many of which are within 100 kb of positional candidate genes. Univariate envGWAS identified 23, 17, and 10 variants associated with temperature, precipitation, and elevation, respectively (Fig S7). The most significant multivariate association in Red Angus is located on chromosome 29 within *BBS1* (Bardet-Biedl syndrome 1), which is involved in energy homeostasis [61]. *BBS1* mutant knock-in mice show irregularities in photoreceptors and olfactory sensory cilia [62] functions that are likely important to an individual’s ability to sense its local environment. This region was not significantly associated in any of the univariate analyses of environmental variables, and was not identified in any of the discrete ecoregion envGWAS. Of the other positional candidate genes identified in this Red Angus analysis, 9 have previously been implicated in adaptive functions in humans or cattle (Table S16). For example, SNPs near *GRIA4* were implicated in body temperature maintenance in cold stressed Siberian cattle [63]. Significant SNPs and their corresponding candidate genes for all three datasets are reported in **Table S15**.

While we found minimal candidate gene overlap between populations, we identified multiple shared biological pathways and processes (**Table S17**) derived from lists of envGWAS positional candidate genes. Pathways in common between populations were driven by largely different gene sets. Across all populations, we identified the “axon guidance” pathway, and numerous other gene ontology (GO) terms related to axon development and guidance as enriched with environmentally-associated loci. Ai et al. (2015) suggested that axon development and migration in the central nervous system is essential for the maintenance of homeostatic temperatures by modulating heat loss or production [64]. In addition to axonal development, a host of other neural signaling pathways were identified in multiple populations. A genome-wide association study for gene-by-environment interactions with production traits in Simmental cattle by Braz et al. (2020) identified a similar set of enriched pathways [28]. These common neural signaling pathways identified by envGWAS are regulators of stress response, temperature homeostasis, and vasoconstriction [65]. We identified other shared pathways involved in the control of vasodilation and vasoconstriction (relaxin signaling, renin secretion, and insulin secretion). Vasodilation and vasoconstriction are essential to physiological temperature control in cattle and other species [66]. The ability to mount a physiological response to temperature stress has a direct impact on cattle performance, making vasodilation a prime candidate for environment-specific selection. Further, vasodilation and vasoconstriction likely also represent adaptation to hypoxic, high elevation environments. Pathways and processes identified by envGWAS signals are reported in **Table S17**.

To further explore the biology underlying adaptive signatures, we performed Tissue Set Enrichment Analysis of our envGWAS candidate gene lists. These analyses using expression data from humans and distantly related worms (*C. elegans*) both identified brain and nerve tissues as the lone tissues where envGWAS candidate genes show significantly enriched expression (**Tables S18-S21**). Tissue-specific expression in the brain further supports our observed enrichment of local adaptation pathways involved in neural signaling and development.

### Identifying loci undergoing region-specific selection with GPSM ecoregion meta-analysis

envGWAS detects allelic associations with continuous and discrete environmental variables, but does not address whether selection is towards increased local adaptation, or whether local adaptation is being eroded by the exchange of germplasm between ecoregions via artificial insemination. We used the spatiotemporal stratification of genotyped animals to identify loci undergoing ecoregion-specific selection. We performed GPSM within each sufficiently genotyped ecoregion and identified variants with high effect size heterogeneity (Cochran’s Q statistic) between ecoregions. Variants with significant heterogeneity across regions that were also significant in at least one within-region GPSM analysis implied ecoregion-specific allele frequency change. These changes could have been due either to selection for local adaptation (Fig 1e), or locally different allele frequencies moving towards the population mean (Fig 1f). We identified 59, 38, and 46 significant SNPs in Red Angus, Simmental, and Gelbvieh, respectively undergoing ecoregion-specific selection. These represent 15, 21, and 26 genomic loci (> 1 Mb to nearest next significant SNP) (Fig 4a). In most cases, these variants have an effect (posterior probability of an effect: m-value > 0.9) in only one or two ecoregions (Fig 4b). Further, nearly all represent the decay of ecoregion-specific allele frequencies towards the population mean (Fig 4c) as opposed to on-going directional selection for ecoregion specific beneficial adaptations (Fig S10-S12). Adaptive alleles at these loci are being driven in frequency towards the population mean allele frequency (Fig 4c), which is typically a low minor allele frequency.

## Discussion

We leveraged large commercially-generated genomic datasets from three major US beef cattle populations to map polygenic selection and environmental adaptation using novel GWAS applications [67]. Using temporally-stratified genotype data we detected very small selection-driven changes in allele frequency throughout the genome. This is consistent with expectations of polygenic selection acting on a large number of variants with individual small effects. Which phenotypes are being selected and driving the allele frequency changes at particular loci is not definitively known. GPSM is a heuristic model, and as a result the SNP effects are not immediately intuitive to interpret in a population genetic context. That said, it allows us to identify the genomic loci responding to selection, and particularly subtle changes due to polygenic selection. GPSM is agnostic to the selected phenotypes, and identifies important loci changing in frequency due to selection without the need to measure potentially difficult or expensive phenotypes. Our GPSM model is of course subject to false-positives and other short comings of genome-wide association models. However, in simulations, GPSM effectively differentiates between selection and drift while accounting for confounding effects such as uneven generation sampling, population structure, relatedness, and inbreeding. Population branch statistics require arbitrary definitions of subpopulations in panmictic populations, and allele frequency trajectories assume random mating, which is violated in our data. The ability of GPSM to account for relatedness and inbreeding likely accounts for the disagreement between GPSM and these other methods. Simulations suggest that GPSM has greater power to detect selection when genotyped individuals are uniformly sampled over time. When genotypes originate from individuals in only the most recent generations the power to detect “old” selection is lessened, and GPSM signatures are enriched for very recent, ongoing selection. As a result, we expect that applying a GPSM-like approach to experimental evolution studies would generate even clearer associations than observed in this study. With the availability of large samples our analytical framework can help solve the long-standing population genetics problem of identifying the loci subjected to polygenic selection.

Future studies exploring the effects of selection from the context of complex trait networks could explain how hundreds or thousands of selected genes act together to shape genomic diversity under directional selection. Candidate genes identified by GPSM suggest selection on pathways and processes involved in production efficiency (growth, digestion, muscle development, and fertility). In addition to a small number of loci, for which function is known, we identify hundreds of novel signatures of ongoing selection.

The envGWAS identified 174, 125, and 130 SNPs associated with both continuous or discrete environmental factors in Red Angus, Simmental, and Gelbvieh, respectively. Using genes found near these environmentally-associated SNPs, we identified a consistent enrichment of pathways and tissues involved in neural development and signaling. These envGWAS associations emphasize the role that the nervous system likely plays in recognizing and responding to environmental stress in mammals, which will be valuable as society and agriculture cope with climate change. In addition to neural pathways, we observe significantly enriched expression of envGWAS genes in the brain tissues of humans, mice, and worms. Other pathways associated with environmental adaptation reveal the importance of mechanisms involved in regulating vasoconstriction and vasodilation, both of which are essential for responses to heat, cold, altitude, and toxic fescue stressors in cattle.

The statistical power and wide geographical distribution of the cattle comprising these data highlights that the utilized approaches can be leveraged to understand the genomic basis of adaptation in many other studies and species. While for GSPM there is a clear connection between the model and directional selection, for envGWAS the model is only identifying associations between the environment and SNP genotypes. These associations could be caused by a multitude of factors, one of which is local adaptation. The small allele frequency differences identified by envGWAS are consistent with a polygenic model of local adaptation, likely driven by small changes in gene expression [68]. Further, envGWAS identifies candidate genes (e.g. *KHDRBS2*) and pathways previously implicated as domestication-related [60]. This suggests that these genes may be under natural or balancing selection to cope with environmental stress, and not specifically part of the domestication process. Further, because different genes in the same pathways were detected in the analyses of the different populations, we hypothesize that these pathways influence local adaptation in many mammals and should be studied in other ecological systems. This knowledge will become increasingly valuable as species attempt to adjust to a changing climate.

The balance between artificial and natural selection in domesticated beef cattle is quite precarious. If natural selection is for a stressor which occurs nation-wide and is positively correlated with production traits, natural selection can effectively act in concert with artificial selection. The identification of genes involved in the immune system by the GPSM analysis reflects this interplay between artificial and natural selection; in other words, natural selection could be acting in the background in populations under artificial selection. However, if natural selection is acting at a local, ecoregion scale, then natural selection and artificial selection via additive genetic breeding values are likely to be at odds. Artificial insemination in cattle has allowed the ubiquitous use of males which have been found to be superior when progeny performance has been averaged across US environments. Due to limited gene flow and phenotypic selection which would act on loci with genotype-by-environment effects, local adaptation likely occurred prior to the 1980s. Our results suggest that environmental associations are present in cattle populations, but that the widespread use of artificial insemination resulting in gene flow and selection on breeding values,, has caused US cattle populations to lose ecoregion-specific adaptive variants [69]. We identified 16, 21, and 30 loci undergoing ecoregion-specific selection in Red Angus, Simmental, and Gelbvieh, respectively. In almost every case, selection has driven allele frequencies within an ecoregion back towards the population mean allele frequency (Fig 1F and Fig 4C). In three independent datasets, we identified a single shared environmentally-associated locus near the gene *KHDRBS2*. This locus has been identified as introgressed in yak, and exhibits an irregular allele frequency trajectory which suggests that it may be subject to balancing selection [70]. Though we identified only a single common envGWAS locus, we observed significant overlap in the pathways regulated by candidate genes within the associated loci. This reveals that adaptive networks are complex and that adaptation can be influenced by selection on functional variants within combinations of genes from these networks. As we work to breed more environmentally-adapted cattle, there will be a need for selection tools that incorporate genotype-by-environment interactions to ensure that cattle become increasingly locally adapted.

We further demonstrate that large commercially-generated genomic datasets from domesticated populations can be leveraged to detect polygenic selection [17] and local adaptation signatures [50]. The identification of adaptive loci can assist in selecting and breeding better adapted cattle for a changing climate. Further, both our statistical approaches and biological findings can serve as a blueprint for studying complex selection and adaptation in other agricultural or wild species. Our results suggest that neural signaling and development are essential components of mammalian adaptation, meriting further functional genomic study. Finally, we observe that local adaptation is declining in cattle populations, which will need to be preserved to sustainably produce protein in changing climates.

## Materials and Methods

RMarkdown files, snakemake files, and R scripts used in the manuscript are available at https://github.com/troyrowan/gpsm_envgwas.

### Genotype data

SNP assays for three populations of genotyped *Bos taurus* beef cattle ranging in density from ∼25K SNPs to ∼770K SNPs were imputed to a common set of 830K SNPs using the large multi-breed imputation reference panel described by Rowan et al. 2019 [71]. Genomic coordinates for each SNP were from the ARS-UCD1.2 reference genome [72]. Genotype filtering for quality control was performed in PLINK (v1.9) [73], reference-based phasing was performed with Eagle (v2.4) [74], and imputation with Minimac3 (v2.0.1) [75]. Following imputation, all three datasets contained 836,118 autosomal SNP variants. All downstream analyses used only variants with minor allele frequencies > 0.01. Upon filtering, we performed a principal component analysis for each population in PLINK. This was to assess if there were discrete subpopulations within the populations and if there were patterns of structure related to ecoregions.

### Generation Proxy Selection Mapping (GPSM)

To identify alleles that had changed in frequency over time, we fit a univariate genome-wide linear mixed model (LMM) using GEMMA (Version 0.98.1) [76]. Here, we used the model:

EQUATION 1:

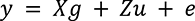

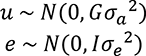

where **y** is an individual’s generation proxy, in our case birth date, and **X** was an incidence matrix that related SNPs to birth dates within each individual and **g** was the estimated effect size for each SNP. An animal’s age as of April 5, 2017 was used as the generation proxy in GPSM. We control for confounding population structure, relatedness, and inbreeding with a polygenic term **u** that uses a standardized genomic relationship matrix (GRM) **G** [20] and we estimated *σ*_*a*_*^2^ and σ*_*e*_*^2^* using restricted maximum likelihood estimation (REML). Here, continuous age served as a proxy for generation number from the beginning of the pedigree. Other than the tested SNP effects, no fixed effects other than the overall mean were included in the model. We tested each SNP for an association with continuous age. To control for multiple-testing, we converted p-values to FDR corrected q-values [77] and used a significance threshold of q < 0.1 to classify significant SNPs. We performed additional negative-control analyses in each dataset by permuting the date of birth associated with each animal’s genotypes to ensure that the detected GPSM signals were likely to be true positives. Permutation was performed ten times for each population. To visualize the allele frequency history of loci undergoing the strongest selection, we fit a loess and simple linear regressions for date of birth and allele frequencies scored as 0, 0.5 or 1.0 within each individual using R [78]. Results were visualized using ggplot2 [79].

### Simulations and gene drops

All simulations were performed in AlphaSimR [80]. Stochastic simulations were performed in 10 replicate sets using 10 sets of founder haplotypes as starting points. We generated founder haplotypes using the AlphaSimR wrapper around MaCS [81]. Using an approximation of the demographic history of cattle, we simulated 10 chromosomes with 20,000 segregating sites each for 2,000 founder individuals (1,000 males and 1,000 females). This resulted in a starting effective population size (N_e_) of approximately 100, similar to estimates of U.S. beef cattle populations [17]. To test other N_e_, we simulated populations with effective population sizes of 50 and 250. Based on the chosen genomic architecture, 1000, 500, or 200 purely additive QTL were randomly assigned to segregating sites. Effect sizes for simulated QTL were drawn from either a normal (mean = 0, variance = 1) or a gamma (shape = 0.42) distribution [82]. Prior to the two divergent selection regimes, we performed five generations of burn-in selection to establish LD in our populations.

After burn-in (generation 0), we performed selection of parents for the next generation in two parallel manners: randomly or using truncation selection on true breeding value. In each scenario, we held the effective population size by selecting appropriate numbers of males and females to be parents each generation. Selection intensity was altered by increasing or decreasing the number of crosses performed (1000, 2000, 4000, 8000). We also varied the number of generations of selection post-burn-in (20, 10, and 5 generations). For each scenario, we extracted 10,000 total simulated individuals for analysis in GPSM. To test the effects of uneven generation sampling that we see in real data, we performed two different strategies for sampling simulated genomes. In one case, we sample an equal number of individuals each generation. In the other, we sample more animals from the most recent generations. The number of sampled individuals is based on a negative exponential distribution that approximates the of ages observed in our real datasets (Fig S1). Sampled individuals were chosen at random, and were not more or less likely to become parents in the next generation. In addition to sampling genotypes each generation, we calculated the allele frequency of simulated QTL each generation to track observed allele frequency changes over the course of selection. This process was performed in replicates of 10 for each scenario, allowing us to calculate descriptive statistics and compare GPSM’s performance across scenarios.

We simulated haplotypes in MaCS for our 5,223 founder individuals. Founder haplotypes spanned 10 chromosomes, each with 20,000 segregating sites for a total of 200,000 SNPs. These founder haplotypes were then randomly dropped through the Red Angus pedigree, restricted to ancestors of genotyped individuals, in a Mendelian fashion, with recombinations occurring at a rate of one crossover per Mb.

After sampling genotypes, we created a standardized genomic relationship matrix (GRM) in GEMMA (v0.98.1) with all SNPs that had a MAF > 0.01. Using GEMMA, we fit the individuals’ true generation number as the dependent variable in a genome-wide linear mixed model. Outputs from GPSM were read, manipulated, and plotted in R using multiple tidyverse packages [83].

Finally, we fit regression models to assess the impact of various parameters on the PVE. For PVE, two regression models were analyzed. In the first, PVE from the random mating simulations was used as the dependent variable with proportion false positives, number of generations, number of crosses, number of QTL, QTL distribution, and number of segregating sites as explanatory variables. In the second, PVE from selection on true breeding value (TBV) was the dependent variable with proportion true positives, number of generations, number of crosses, number of QTL, QTL distribution, and number of segregating sites as dependent variables.

### Birth date variance component analysis

To estimate the amount of variation in birth date explained by GPSM significant SNPs, we performed multi-GRM GREML analyses for birth date in GCTA (v1.92.4) [84]. We built separate GRMs using genome-wide significant markers and all remaining makers outside of significant GPSM loci (> 1 Mb from significant GPSM SNPs to control for markers physically linked to significant GPSM SNPs). To further partition the variance in birth date explained by subsets of SNPs, we performed a GREML analysis using three GRMs created with genome-wide significant (q < 0.1) SNPs, an equal number of the next most significant SNPs, and an equal number of randomly selected markers not present in the first two classes with minor allele frequencies > 0.15, to match the allele frequencies of significant SNPs. These three GRM were each constructed using 268, 548, and 763 SNPs for Red Angus, Simmental, and Gelbvieh, respectively.

### Allele frequency time series methods for detecting selection

We used the software wfABC (Wright-Fisher Approximate Bayesian Computation) [33] to generate estimates of the selection coefficient at each locus. We partitioned individuals into generations based on the maximum generation number estimated from the Red Angus pedigree using the optiSel R package [85]. Since wfABC assumes random-mating populations of unrelated individuals, we also performed relationship pruning in PLINK, removing individuals with estimated relationship coefficients > 0.0625. The wfABC software estimated selection coefficients for each imputed SNP with MAF > 0.01, using an effective population size of 150 [17].

### TreeSelect

Using TreeSelect [86], we artificially subdivided our genotyped populations to compare statistics from a population branch statistic (PBS) method with GPSM effect sizes and p-values. TreeSelect tests for allele frequency differences between discretely labeled, but closely related populations. We ran TreeSelect two ways: First, where each genotyped population (Red Angus, Simmental, and Gelbvieh) was its own branch, and then in three separate analyses where we subdivided each population into three equally sized groups based on an individual’s birth date. These three age classes consisted of the oldest ⅓ of individuals, the middle ⅓ of the age distribution, and the youngest ⅓ of individuals. We compared TreeSelect chi-squared values with GPSM betas to quantify overall relationships between the statistics.

### Environmental data

Thirty-year normals (1981-2010) for mean temperature ((average daily high (°C) + average daily low (°C)/2), precipitation (mm/year), and elevation (m above sea level) for each 4 km^2^ of the continental US were extracted from the PRISM Climate Dataset [87], and used as continuous dependent variables in envGWAS analysis. Optimal *K*-means clustering of these three variables grouped each 4 km^2^ of the continental US into 9 distinct ecoregions. Using the reported breeder zip code for each individual, we linked continuous environmental variables to animals and partitioned them into discrete environmental cohorts for downstream analysis. For ecoregion assignments, latitude and longitude were rounded to the nearest 0.1 degrees. As a result, some zip codes were assigned to multiple ecoregions. Animals from these zip codes were excluded from the discrete region envGWAS but remained in analyses that used continuous measures as dependent variables.

### Environmental Genome-wide Association Studies (envGWAS)

To identify loci segregating at different frequencies within discrete ecoregions or along continuous climate gradients, we used longitudinal environmental data for the zip codes attached to our study individuals as dependent variables in univariate and multivariate genome-wide LMMs implemented in GEMMA (Version 0.98.1). We fit three univariate envGWAS models that used 30-year normal temperature, precipitation, and elevation data as dependent variables. These used an identical model to EQUATION 1, but used environmental values as the dependent variable (**y**) instead of birth date. We also fit a combined multivariate model using all three environmental variables to increase power. To identify loci associated with entire climates as opposed to only continuous variables, we fit univariate and multivariate case-control envGWAS analyses using an individual’s region assignment described in the “Environmental Data” section as binary phenotypes. Proportion of variation explained (PVE), phenotypic correlations, and genetic correlations were estimated for continuous environmental variables and discrete environmental regions using GEMMA’s implementation of REML.

To ensure that envGWAS signals were not driven by spurious associations, we performed two separate permutation analyses. In the first, we randomly permuted the environmental variables and regions associated with an individual prior to performing each envGWAS analysis, detaching the relationship between an individual’s genotype and their environment. In the second, to ensure that envGWAS signals were not driven by the over-sampling of individuals at particular zip codes, we permuted the environmental variables associated with each zip code prior to envGWAS analysis. These two types of permutation analyses were performed for each dataset and for each type of univariate and multivariate envGWAS analysis. We determined significance using a permutation-derived p-value cutoff (p < 1⨯10^-5^) [88].

### GPSM meta-analyses

To identify variants undergoing ecoregion-specific allele frequency changes, we performed GPSM analyses within each region with more than 600 individuals. The SNP significance testing effects and standard errors from each of the within-region GPSM analyses were combined into a single meta-analysis for each population using METASOFT (v2.0.1) [89]. We identified loci with high heterogeneity in allele effect size, suggesting region-specific selection. An m-value indicating the posterior-probability of a locus having an effect in a particular ecoregion was calculated for each of these loci [90].

### Gene set and tissue set enrichment analysis

Using the NCBI annotations for the ARS-UCD1.2 *Bos taurus* reference assembly, we located proximal candidate genes near significant SNPs from each of our analyses. We generated two candidate gene lists each from significant GPSM and envGWAS SNPs. Lists contained all annotated genes within 10 kb from significant SNPs. We consolidated significant SNPs from all envGWAS analyses to generate a single candidate gene list for each breed. Using these candidate gene lists, we performed gene ontology (GO) and KEGG pathway enrichment analysis using Clue GO (v2.5.5) [91] implemented in Cytoscape (v3.7.2) [92]. We identified pathways and GO terms where at least two members of our candidate gene list comprised at least 1.5% of the term’s total genes. We applied a Benjamini-Hochberg multiple-testing correction to reported p-values and GO terms with FDR corrected p-values < 0.1 were considered significant.

Using the above gene sets, we performed three separate Tissue Set Enrichment Analyses (TSEA) using existing databases of human, mouse, and worm gene expression data. We searched for enriched gene expression with data from the Human Protein Atlas [93] and Mouse ENCODE [94] using the Tissue Enrich tool (v1.0.7) [95]. Additionally, we performed another Tissue Set Enrichment Analysis using GTEx data [96] and a targeted Brain Tissue Set Enrichment Analysis in the pSI R package (v1.1) [97]. Finally, we used Ortholist2 [98] to identify *C. elegans* genes orthologous with members of our envGWAS and GPSM gene lists. We then queried these lists in WormBase’s Tissue Enrichment Analysis tool [99, 100] to identify specific tissues and neurons with enriched expression in *C. elegans*. We used each tool’s respective multiple-testing correction to determine significance. We deemed an enrichment in a tissue “suggestive” when its p-value was < 0.1.

### Summary Data

Summary data from GPSM and envGWAS analyses are publicly available as a Zenodo repository [125].

## Acknowledgments

We appreciate comments from Wes Warren, Jeremy Taylor, and William Lamberson while writing this manuscript and advice from Iain Mathieson on allele trajectory analyses. We also appreciate farmers and ranchers for collecting this data and for data being shared from the Red Angus Association of America, American Simmental Association, and American Gelbvieh Association.

**S1 Supplementary Information for Powerful detection of polygenic selection and environmental adaptation in US beef cattle.** This file includes supplementary text, Figures S1 to S11, Tables S1, S3, S4, S6, S7, S11-S14, S16, and SI References.

**S2 Large Supplementary Tables.** This file includes supplementary Tables S2, S5, S8, S9, S10, S15, S17, S18, S19, S20 and S21. Legends for these tables are included in S1 Supplementary Information.

## Supplementary Information

### Supplementary Text

#### GPSM detects signatures of polygenic selection across and within three populations of U.S. Beef Cattle

In addition to the locus undergoing a massive allele frequency shift on BTA28 (lead SNP *rs1762920*), we identify 6 other loci under selection in all three populations. (Table S7). These shared GPSM signals suggest that not only are there similar selection pressures, but common genomic architectures under selection in these three populations. While we identify significantly more population-specific GPSM signatures, shared signatures are of interest as they are likely serving an important role in all breeds of beef cattle. We discuss these loci and their potential functions in beef cattle that are driving allele frequency changes.

The largest genomic region detected by GPSM lies at the end of BTA1 (157.5 Mb - 158.5 Mb). The lead SNP in this peak (*rs1755753*) lies within the long non-coding RNA (lncRNA) *LOC112448253*. This lncRNA has not been previously associated with any traits or function in cattle. This region contains dozens of other potential candidate genes. This selected region is also immediately upstream of *PRDM9*, a modulator of recombination in most mammalian species, including cattle [1–3]. While no variants within PRDM9 reach genome-wide significance, variants ∼10.7 kb from the TSS are responding to selection. Selection on *PRDM9* could increase average recombination rate or allow novel motif binding in order to create novel favorable haplotype combinations [4].

We identified six other common genomic regions under strong selection in all three populations, encompassing 106 statistically significant markers. Another 90 SNPs overlap in at least two populations, corresponding to 15 additional genomic regions under selection (Table S7). All three populations have been selected for increased growth traits over the last 50 years [5], and GPSM identifies two genomic regions that have been previously associated with feed efficiency and growth traits. The common peak on BTA12 (lead SNP *rs1389713*) is ∼182 kb upstream of the *DACH1* (Dachshund homolog 1), a transcription factor associated with post-weaning gain, various indicators of feed efficiency [6, 7], and backfat thickness [8] in cattle. The shared GPSM peak on BTA14 resides near a known QTL for post-weaning gain near the gene *LRP12* (LDL receptor related protein 12) [9].

In addition to selection on loci that appear directly involved in growth and efficiency, we identify multiple selection targets likely involved in aspects of immune function. A shared significant peak on BTA2 (lead SNP *rs1080110*) resides within the *ARHGAP15* gene (Rho GTPase Activating Protein 15) that is essential for Trypanosomiasis resistance in African cattle populations [10–12]. Though Trypanosomiasis and other tsetse fly-transmitted diseases are restricted to Africa, genetic tolerance to similar immune disturbances may account for the positive selection observed in these three American cattle populations. Variants within *ADORA1* (Adenosine A1 Receptor) are also detected as being under selection by GPSM. In cattle, *ADORA1* plays a role in the activation of polymorphonuclear neutrophilic leukocytes, which are important for peripartial immune responses in cattle [13], and likely play roles in other immune functions. *ADORA1* and other purinergic receptors play an important role in bone metabolism [14] and likely growth in cattle. This signature within *ADORA1* also spans a potentially regulatory region for *MYBPH*, an important gene in muscle formation and development, and another potential target of selection [15]. In this case and others, we identify multiple logical candidate genes within regions undergoing selection. We report a complete listing of significant SNPs and candidate genes from GPSM analyses of each population in (**Table S5**).

We observe 22 genomic regions that are changing in frequency in at least two of our datasets. In addition to these shared loci, we identify evidence of shared networks and genetic architectures under selection (**Table S8**). To discern common biological pathways and processes under selection across multiple populations, we identified genes within 10 kb of SNPs identified by GPSM in at least two populations and performed a gene enrichment analysis in ClueGO. Using the 46 genes residing in or near the 29 shared GPSM signatures identified by at least two datasets identified multiple biological processes undergoing selection. The most significant pathways involved G-protein coupled signaling (Benjamini–Hochberg adjusted p-value = 0.038) and purinergic receptor signaling pathways (Benjamini–Hochberg adjusted p-value = 0.034, associated genes *ADORA1* and *P2RY8*). Purinergic receptors have been identified as important drivers of immune responses in cattle [13]. Selection on immune pathways is likely driven by the increased production efficiency of healthy calves [16, 17]. Shared selection on *SLC2A5* and *BIRC5* point towards biological pathways involved in sensing carbohydrate, hexose, and monosaccharide stimuli (Benjamini-Hochberg adjusted p-value = 0.01). We also expect that an enhanced metabolic response to carbohydrates would result in increased animal efficiency. Finally, genes involved in the regulation of arterial blood pressure (Benjamini–Hochberg adjusted p-value = 0.045: *ADORA1*, *SLC2A5*) made up the lone other significant gene class under selection across all three populations. Gene enrichment analysis within populations also identified population-specific pathways and processes under selection. In Simmental cattle we detect 23 GPSM candidate genes involved in olfactory transduction. While not related directly to growth traits, olfactory receptors have been identified as selection targets in many mammalian species, including cattle [18–20]. This rapidly-evolving class of genes is also associated with growth and carcass traits, suggesting a wide range of functions that are not limited to detecting smell [9, 21].

#### envGWAS identifies adaptive pathways and processes

Local adaptation is likely highly complex and controlled by many areas of the genome. Though we detected minimal overlap in candidate genes across datasets, we identified multiple conserved biological processes and pathways that appear to play roles in local adaptation across populations.

Though there was minimal candidate gene overlap between continuous and discrete envGWAS (19 of 187 total envGWAS candidate genes in Red Angus), we identified many shared pathways and gene ontologies. In most cases where we observed GO or pathway overlap, statistical significance was greater for the discrete envGWAS analysis, simply due to the difference in number of provided genes (168 vs 38). Most of the shared terms and pathways were driven by the genes *BAD*, *EFNA5*, *LRRC4C*, *MARK2*, *PLCB3*, and *PRKG2* detected in both analyses. In these overlapping terms, additional genes from the discrete zone envGWAS further supplemented the identified terms.

We performed gene enrichment analyses with gene lists from all univariate and multivariate, discrete and continuous envGWAS analysis in each population. This allowed sufficiently large gene lists to identify potentially adaptive pathways and processes. We were particularly interested in identifying shared pathways and processes between populations since the number of shared genomic regions was low. Across all three populations, we consistently identified the “axon guidance” pathway, and numerous GO terms relating to axon development and guidance under region-specific selection. Ai et al. (2015) [22] suggested that axon development and migration in the central nervous system is essential for the maintenance of homeostatic temperatures by modulating heat loss or production [23]. The direction and organization of axons is an essential component of the olfactory system which is frequently implicated in environmental adaptation through the recognition of local environmental cues [24, 25]. Other pathways identified across all three datasets include “cholinergic synapse”, “glutamatergic synapse”, and “platelet activation”. Cholinergic signaling drives cutaneous vascular responses to heat stress in humans [26–28]. Additionally, cholinergic receptors act as the major neural driver of sweating in humans [29]. Glutamatergic synapses are involved in neural vasoconstriction [30, 31]. “Retrograde endocannabinoid signaling”, “dopaminergic synapse”, “GABAergic synapse”, and “serotonergic synapse” pathways are also significantly enriched by envGWAS candidate genes. Nearly all of these important neural signaling pathways are also enriched in a series of gene-by-environment interaction GWAS for birth weight, weaning weight, and yearling weight in Simmental cattle by [32]. Taken together, these results suggest that pathways involved in neuron development and neurotransmission are essential components of local adaptation in cattle. Temperature homeostasis is largely controlled by the central nervous system [33], making environment-specific selection on these pathways an efficient way for populations to adapt.

Other pathways identified by envGWAS appear to play important roles in vasodilation and vasoconstriction. Relaxin signaling was identified in both Red Angus and Simmental populations as a locally adaptive pathway under selection. Relaxin, initially identified as a pregnancy-related hormone, is an important modulator of vasodilation [34]. In Simmental this pathway association originates from local adaptation signatures near the genes *COL1A1*, *MAPK10*, and *PRKACA* identified both in our discrete multivariate envGWAS and in the Desert ecoregion univariate analysis. Relaxin signaling was also identified in Red Angus, but with four entirely different genes (*LOC529425*, *PLCB3*, *PRKACB*, *VEGFB*). While all four of the Red Angus genes were identified in multivariate analyses, we identify two of them in the Desert ecoregion univariate envGWAS. Vasodilation is an essential component of physiological temperature adaptation in cattle and other species [35–37]. The ability to mount a physiological response to heat stress has a direct impact on cattle performance. Heat stressed cattle have decreased feed intake, slower growth rates, and decreased fertility [38]. The Renin secretion pathway, which is also directly involved in vasoconstriction was identified in Red Angus (*ADRB1*, *PLCB3*, *PRKACB*, *PRKG2*), and has also been previously implicated in physiological responses to heat stress in cattle [39]. In each population we identify multiple biological processes related to the regulation of insulin secretion. Insulin secretion is elevated in heat stressed cattle [40] and pigs [41], suggesting that it plays a role in metabolism and thermoregulation. Insulin secretion in response to the presence of glucose may also be related to different diets and forage availability along these continuous environmental gradients [42].

Other pathways identified by envGWAS candidate genes from Simmental point towards the immune system’s role in local adaptation. “Th1 and Th2 cell differentiation” and “Th17 cell differentiation” were significant in a KEGG pathway analysis. The development of these cell types is essential for adaptive immune responses [43, 44]. These signals were driven in part by region-specific allele frequency differences in or near *MAPK10* an innate immunity gene identified in human studies as an adaptive target of selection [45]. We also identify multiple cardiac-related pathways from Simmental envGWAS genes (“Dilated cardiomyopathy”, “Arrhythmogenic right ventricular cardiomyopathy”, “Hypertrophic cardiomyopathy”) driven by a pair of related genes *SGCA* and *SGCD*. Like other circulatory-related pathways, the alleles driving this signal were identified in both multivariate and Desert ecoregion envGWAS. We expect that these cardiac-related pathways, like renin secretion and relaxin signaling affect the efficiency of circulation and improved temperature homeostasis when exposed to heat or cold stress. Selection on cardiovascular function is also likely a central component of adaptation to high altitude [46].

#### Tissue Set Enrichment Analysis

To further disentangle the biological basis of GPSM and envGWAS adaptive signatures, we performed a series of Tissue Set Enrichment Analyses (TSEA) based on gene expression data from humans and worms (*C. elegans*). These analyses identified tissues in which our envGWAS candidate genes were preferentially expressed. Our candidate gene lists for each population consisted of annotated cattle genes within 10 kb of a significant GPSM SNPs or SNPs identified in any of our multivariate or univariate envGWAS analysis. In using expression data from other species, non-orthologous cattle genes are not included in enrichment analyses.

Using GPSM candidate genes in TSEA identified enriched tissues that correspond to population-specific production traits known to be under selection. Using gene expression measures from the GTEx pilot dataset [47] in the pSI R package [48] we identify suggestive enriched expression in various reproductive-related tissues from the Red Angus GPSM candidate gene set. We observe suggestive enrichments (p < 0.1) for human breast, ovary, pituitary, and uterus tissues (**Table S9-S10**) among others. An analysis of this gene list with gene expression data from the Human Protein Atlas also identified significantly enriched expression in cervix, uterus, and brain tissues (FDR-corrected p-value < 0.1) and suggestive expression fold change in the ovary. These results provide further evidence that selection on fertility and reproductive traits have been ongoing in the Red Angus population over the last ∼10 generations (Red Angus Association of America EPD trends https://redangus.org/genetics/epd-trends/). Further, using Gelbvieh GPSM candidate gene sets, we identify enriched expression in numerous tissues including skeletal muscle, nerve, thyroid and adipose tissue. These enriched tissues align with known ongoing selection for increased growth and carcass quality. Despite the numerous enriched pathways and processes from the Simmental GPSM gene lists, we identified minimal tissue-specific expression. We did not identify significant or suggestive tissue enrichments in the Simmental GPSM gene set using the Human Protein Atlas in TissueEnrich. Uterine tissue was the only tissue showing suggestive enrichment.

Pathway analyses of envGWAS candidate gene sets in all three populations pointed towards a role in neural development and signaling in modulating adaptation. We used TSEA in humans and *C. elegans* to provide further evidence of brain and nervous system tissues involvement in environmental adaptation. Using *C. elegans* tissues allowed us to refine the expression of conserved genes to individual neuron resolution. Using gene expression data from the Human Protein Atlas in the TissueEnrich software, we identify the cerebral cortex to be the lone significant tissue among our candidate gene sets from Red Angus and Gelbvieh envGWAS analyses (**Table S18-S19**). These results agreed with TSEA using GTEx data. Despite identifying similar neural pathways in the Simmental population, we did not observe enriched expression in brain tissue in either human TSEA with this gene list.

To further probe the specific brain regions expressing envGWAS candidate genes, we performed a brain-specific expression enrichment analysis using the pSI tool with human brain expression data from BrainSpan and the Allen Brain Atlas [49]. Complete results are reported in **Supplementary Tables 20-21.** The only brain region significantly enriched for envGWAS candidate genes was the cortex in Simmental (p = 1.525×10^-4^). Interestingly, Simmental was the only population in which Brain tissue did not show enriched expression in the GTEx data. The envGWAS candidate genes from Red Angus and Gelbvieh showed suggestive enrichments in expression in the Cerebellum (p = 0.096) and Striatum, respectively (p = 0.059). While some suggestive cell-type specific expression differences existed with each gene list, we did not observe any tissues with conserved expression across populations.

---

**Fig S1.**
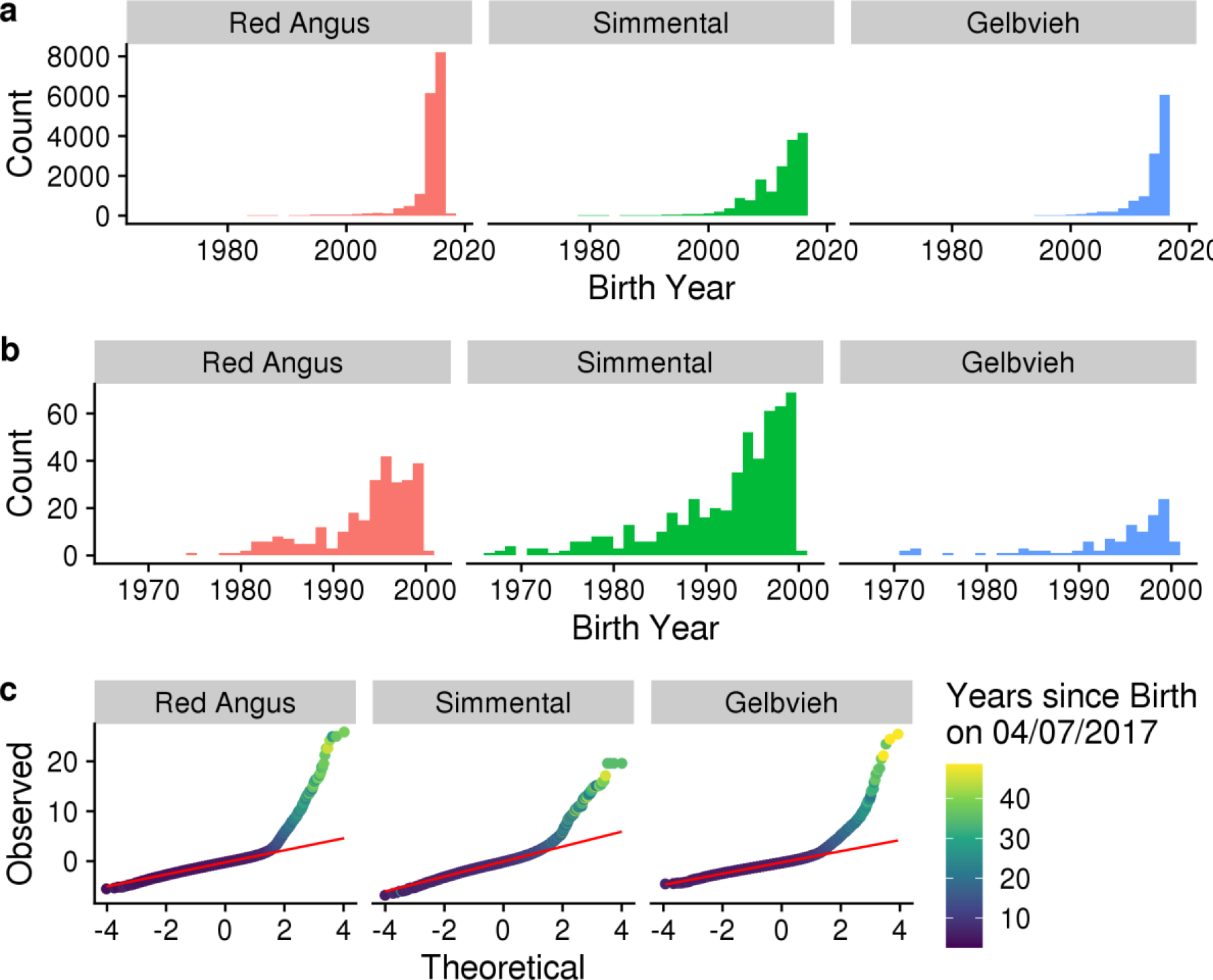
Distributions of continuous birth date in sampled Red Angus, Simmental, and Gelbvieh populations. (a) Birth date histograms for complete datasets. (b) Histograms of animal birth dates born before 2000. (c) Q-Q plots of residual error from a GREML analysis of birth date in each population. Points represent individuals, and are colored by the number of years since the animal’s birth date.

**Fig S2.**
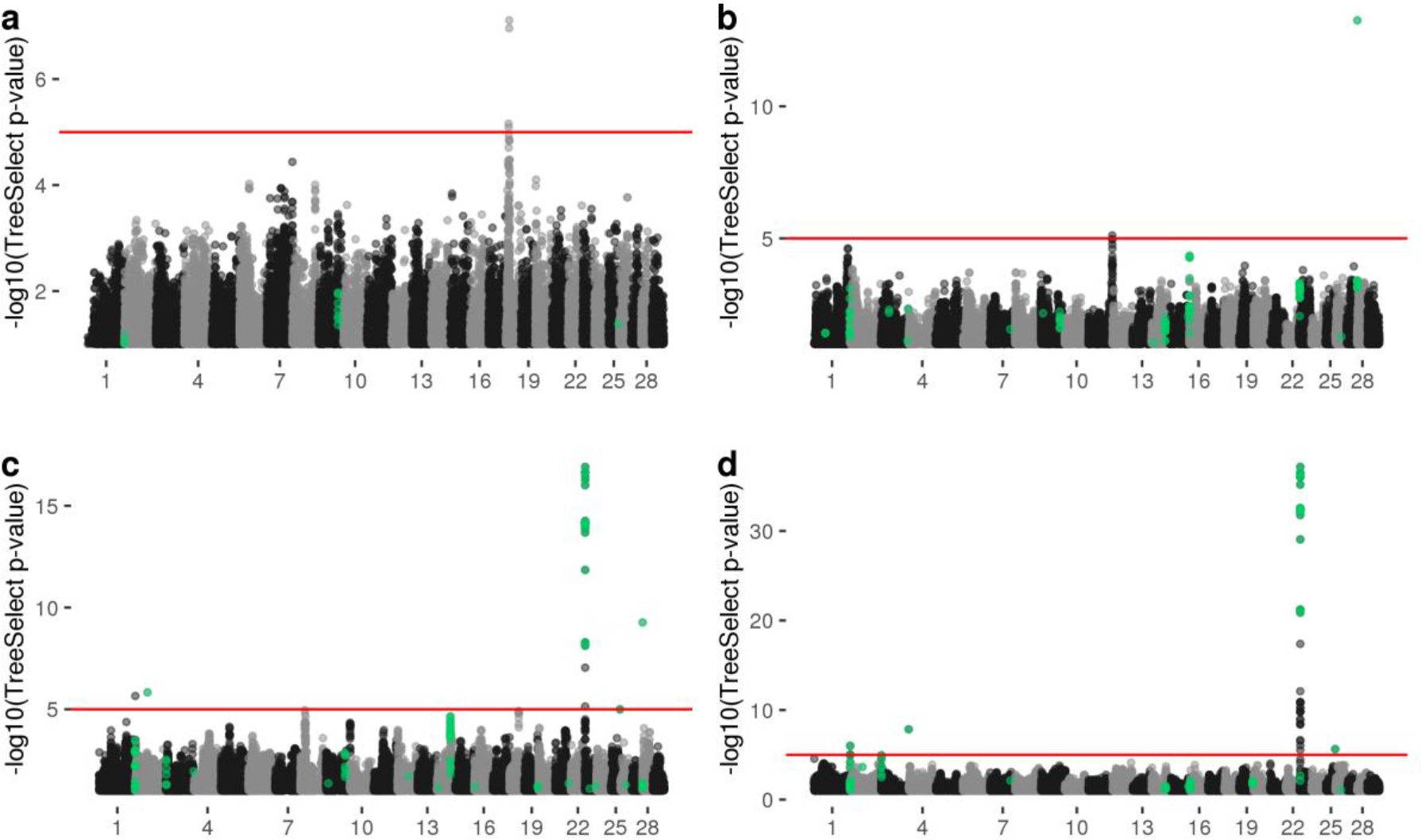
TreeSelect results for Red Angus dataset. a) Single SNP -log_10_(p-values) for Red Angus branch of across-breed TreeSelect analysis. TreeSelect Manhattan plots for a) oldest ⅓, b) middle ⅓, and c) youngest ⅓ branches in within-breed analysis for the Red Angus population. Red line indicates significance at p < 1 x 10^-5^. Green points are SNPs that were significant (q < 0.1) in GPSM analysis of Red Angus dataset.

**Fig S3.**
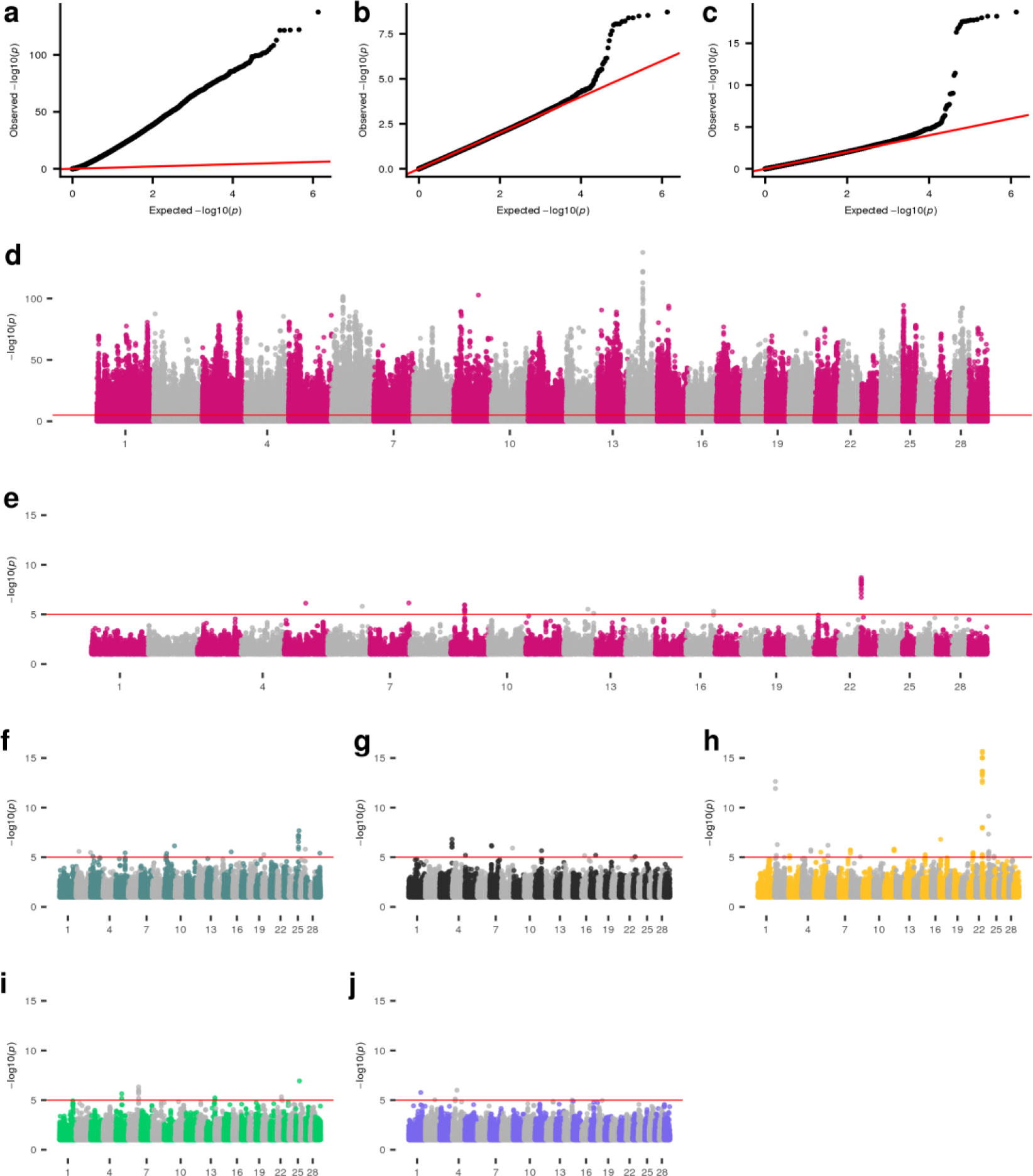
Manhattan plots of discrete envGWAS in Red Angus cattle. Q-Q plots for envGWAS p-values of (a) a linear model for Forested Mountains ecoregion membership, (b) a linear mixed model for Forested Mountain ecoregion membership, and (c) a multivariate linear mixed model of ecoregion membership. Univariate discrete envGWAS for (d) Forested Mountain linear model, (e) Forested Mountains linear mixed model, (f) Southeast, (g) Fescue Belt, (h) Arid Prairie, (i) High Plains, and (j) Upper Midwest & Northeast ecoregions. In all Manhattan plots the red line indicates an empirically-derived p-value significance threshold from permutation testing (p < 1×10-5). Note the drastically inflated p-values from the linear model in (a). Further, note that associated loci are not consistent between linear model and linear mixed model, highlighting the need to control for geographic dependency with a genomic relationship matrix.

**Fig S4.**
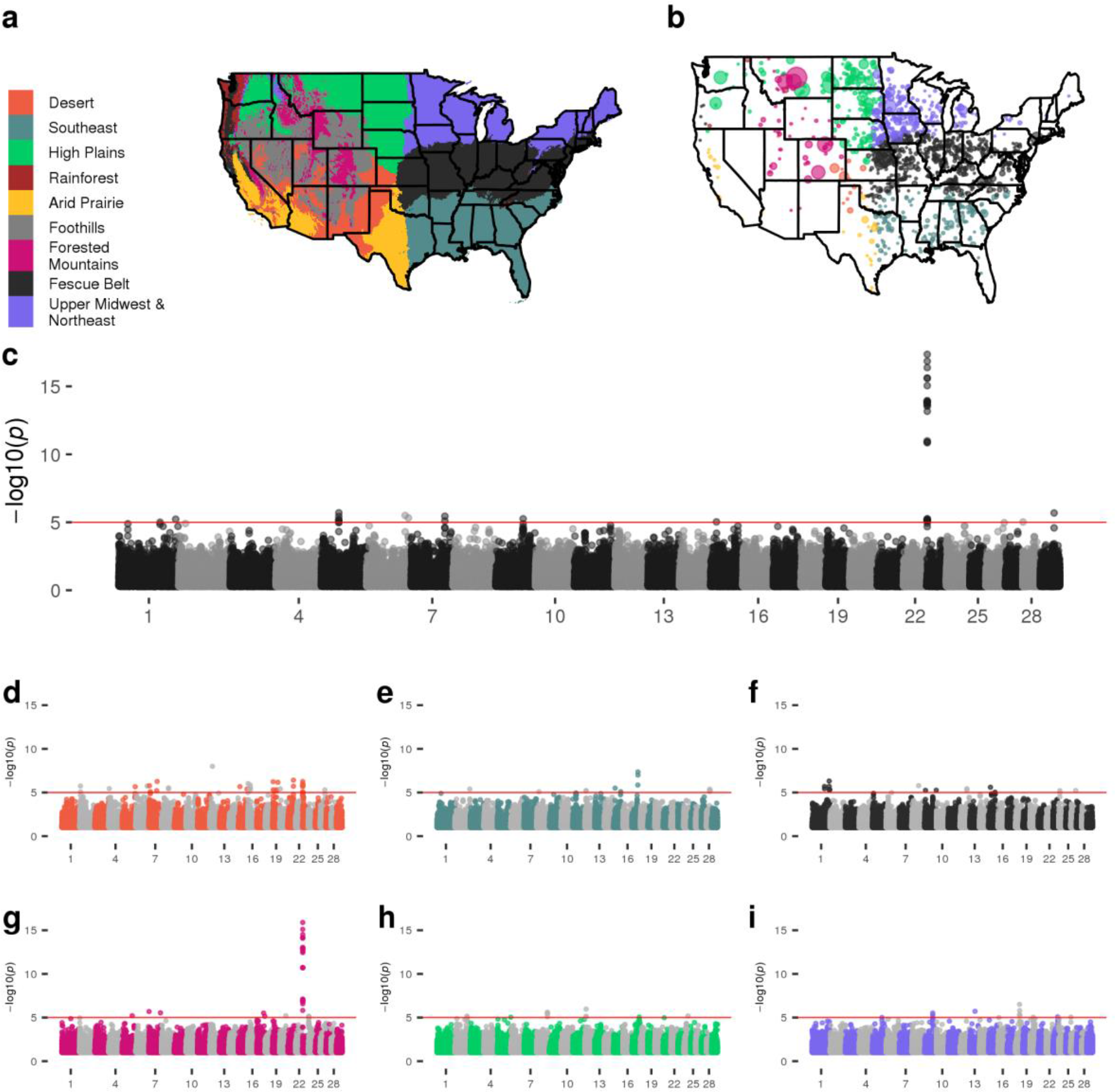
Manhattan plots of discrete envGWAS in Simmental cattle. (a) Nine ecoregions of the continental United States defined by *K*-means clustering of 30-year normal temperature, precipitation, and elevation. (b) Locations of Simmental animals colored by breeder’s ecoregion and sized by number of animals at that location. (c) Multivariate envGWAS (case-control for regions with > 600 animals). Univariate discrete envGWAS for (d) Desert, (e) Southeast, (f) Fescue Belt, (g) Forested Mountains, (h) High Plains, and (i) Upper Midwest & Northeast ecoregions. In all Manhattan plots the red line indicates an empirically-derived p-value significance threshold from permutation testing (p < 1×10^-5^).

**Fig S5.**
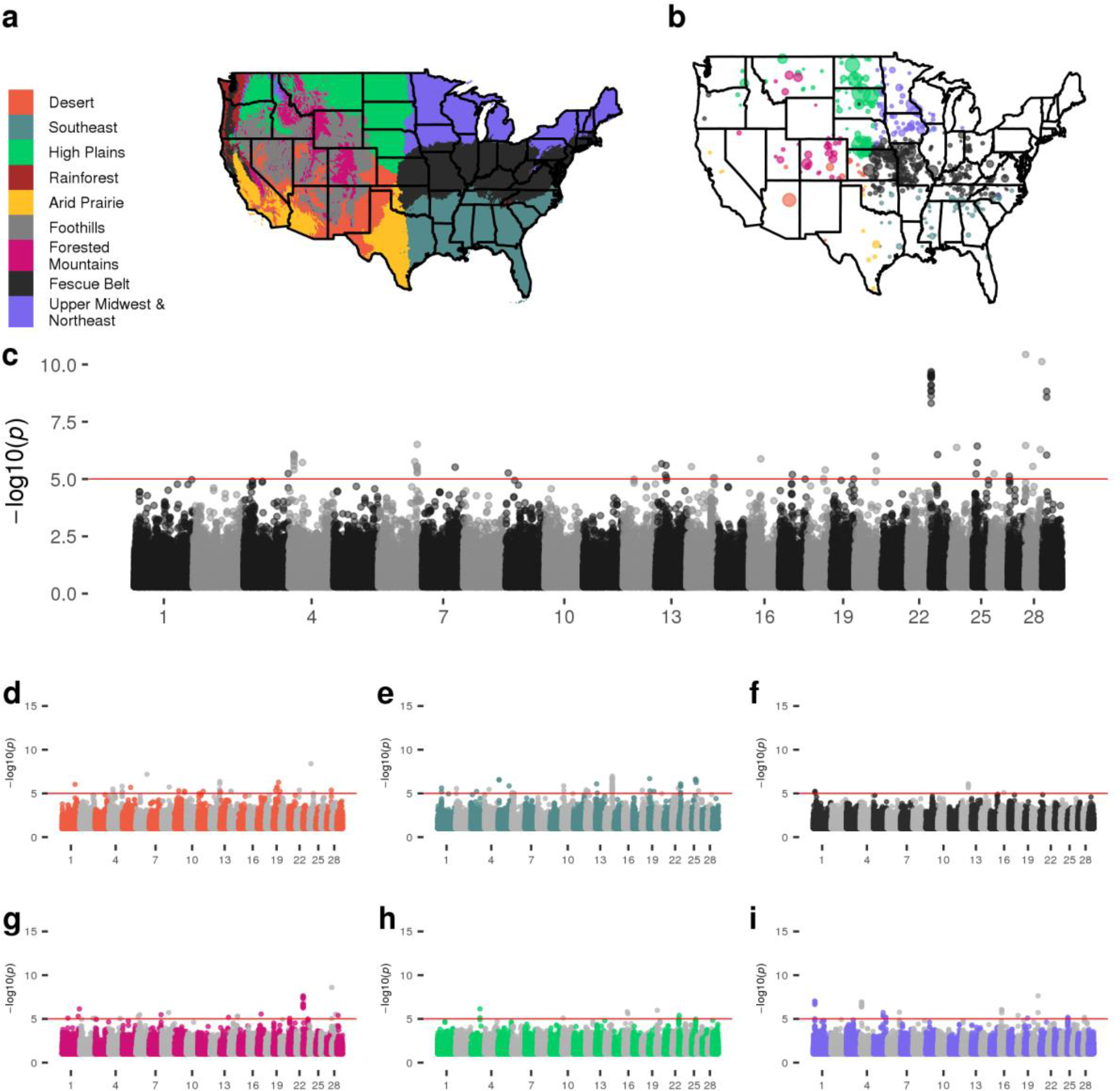
Manhattan plots of discrete envGWAS in Gelbvieh cattle. (a) Nine ecoregions of the continental United States defined by *K*-means clustering of 30-year normal temperature, precipitation, and elevation. (b) Locations of Gelbvieh animals colored by breeder’s ecoregion and sized by number of animals at that location. (c) Multivariate envGWAS (case-control for regions with > 600 animals). Univariate discrete envGWAS for (d) Desert, (e) Southeast, (f) Fescue Belt, (g) Forested Mountains, (h) High Plains, and (i) Upper Midwest & Northeast ecoregions. In all Manhattan plots the red line indicates an empirically-derived p-value significance threshold from permutation testing (p < 1×10^-5^).

**Fig S6.**
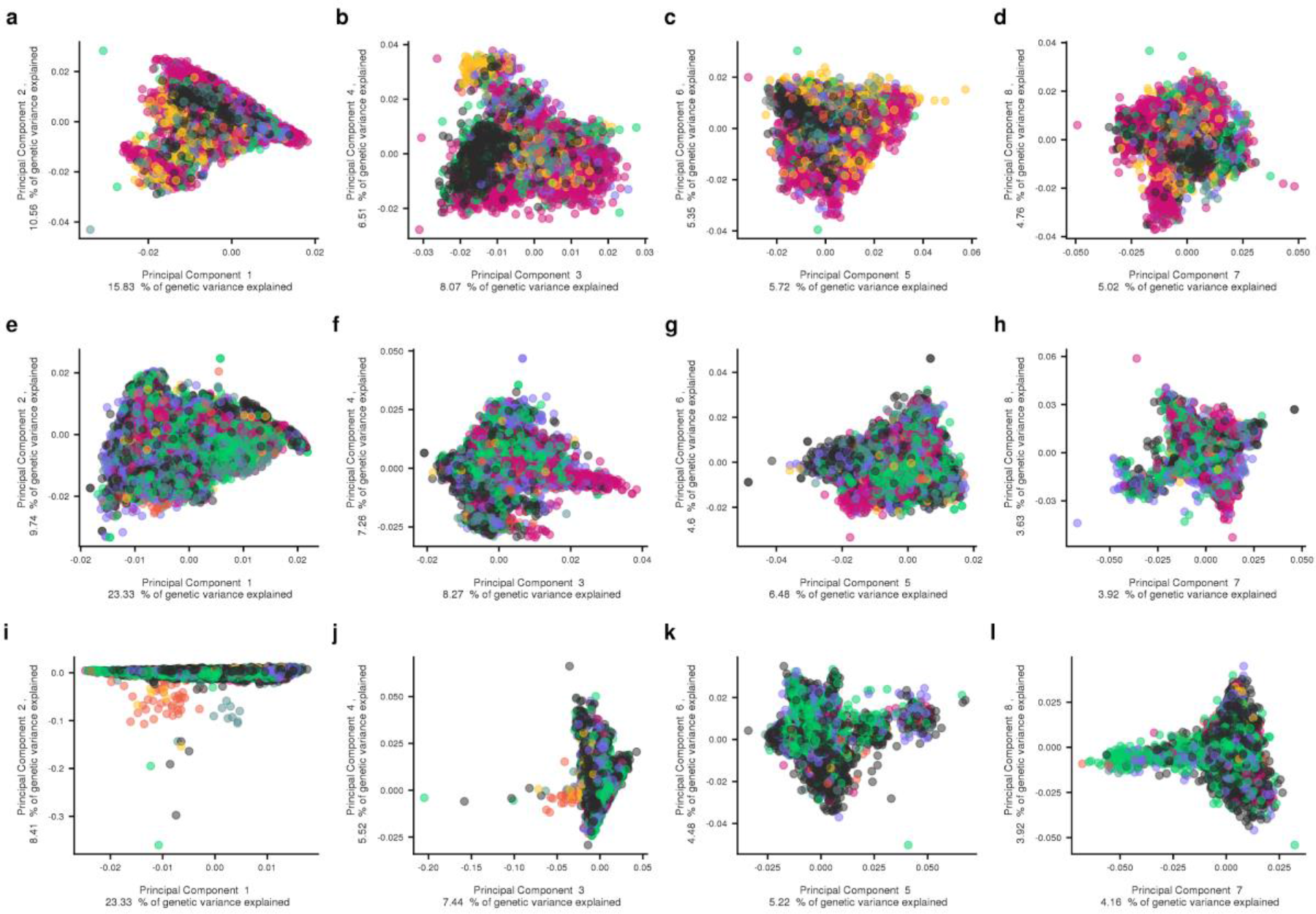
Plots of first eight principal components from PCA analysis. Plots for Red Angus (A-D), Simmental (E-H), and Gelbvieh (I-L). Points indicate individuals, colored by their assigned ecoregion.

**Fig S7.**
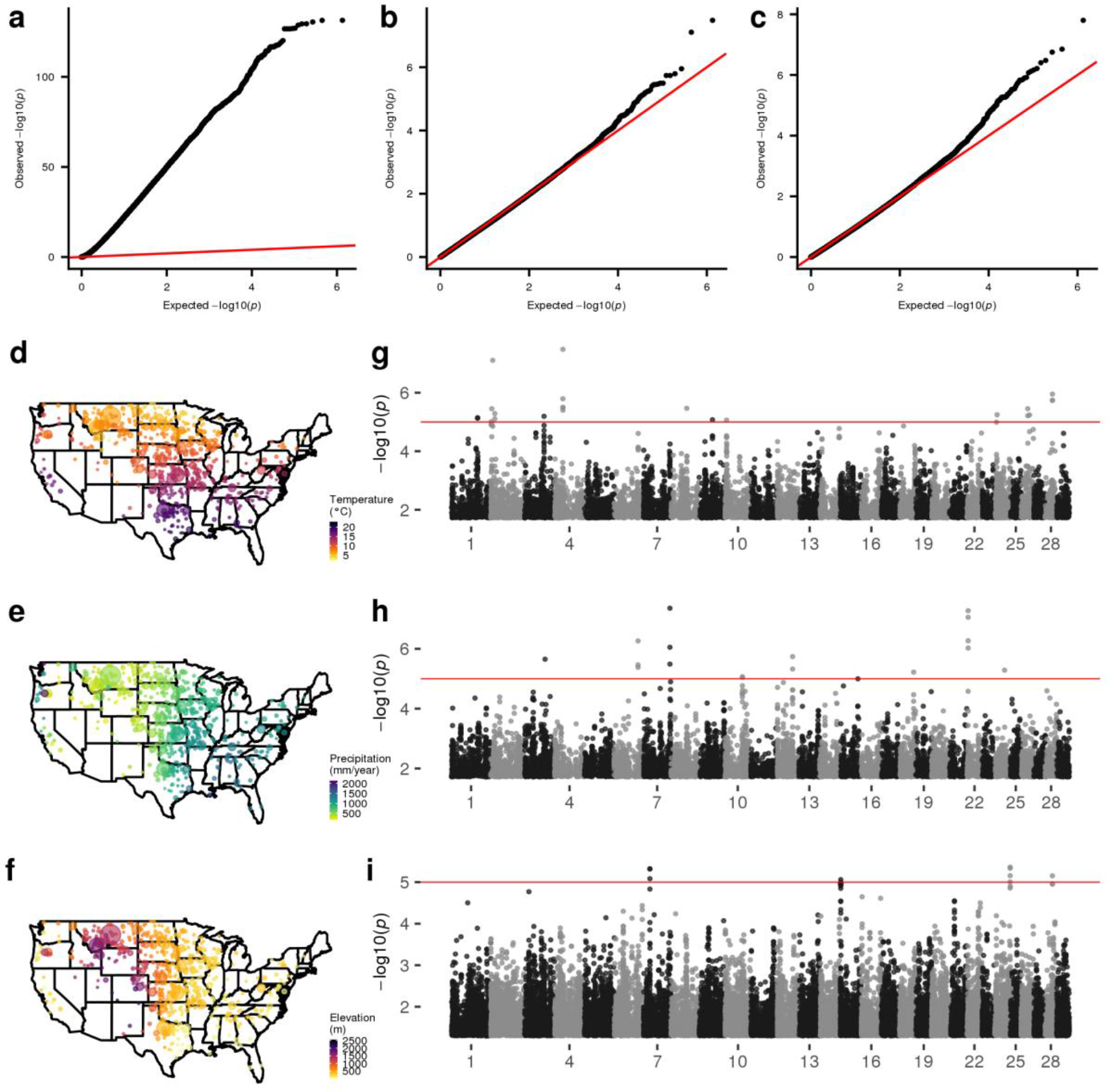
Continuous environmental variable envGWAS in Red Angus cattle. Q-Q plots for envGWAS p-values of (a) a linear model for temperature, (b) a linear mixed model for temperature, and (c) a multivariate linear mixed model of temperature, precipitation, and elevation. Geographic distributions colored by (d) temperature, (e) precipitation, (f) elevation. Manhattan plots for univariate envGWAS analysis of (g) temperature, (h) precipitation, (i) elevation. Red lines indicate permutation-derived p-value cutoff of 1×10^-5^.

**Fig S8.**
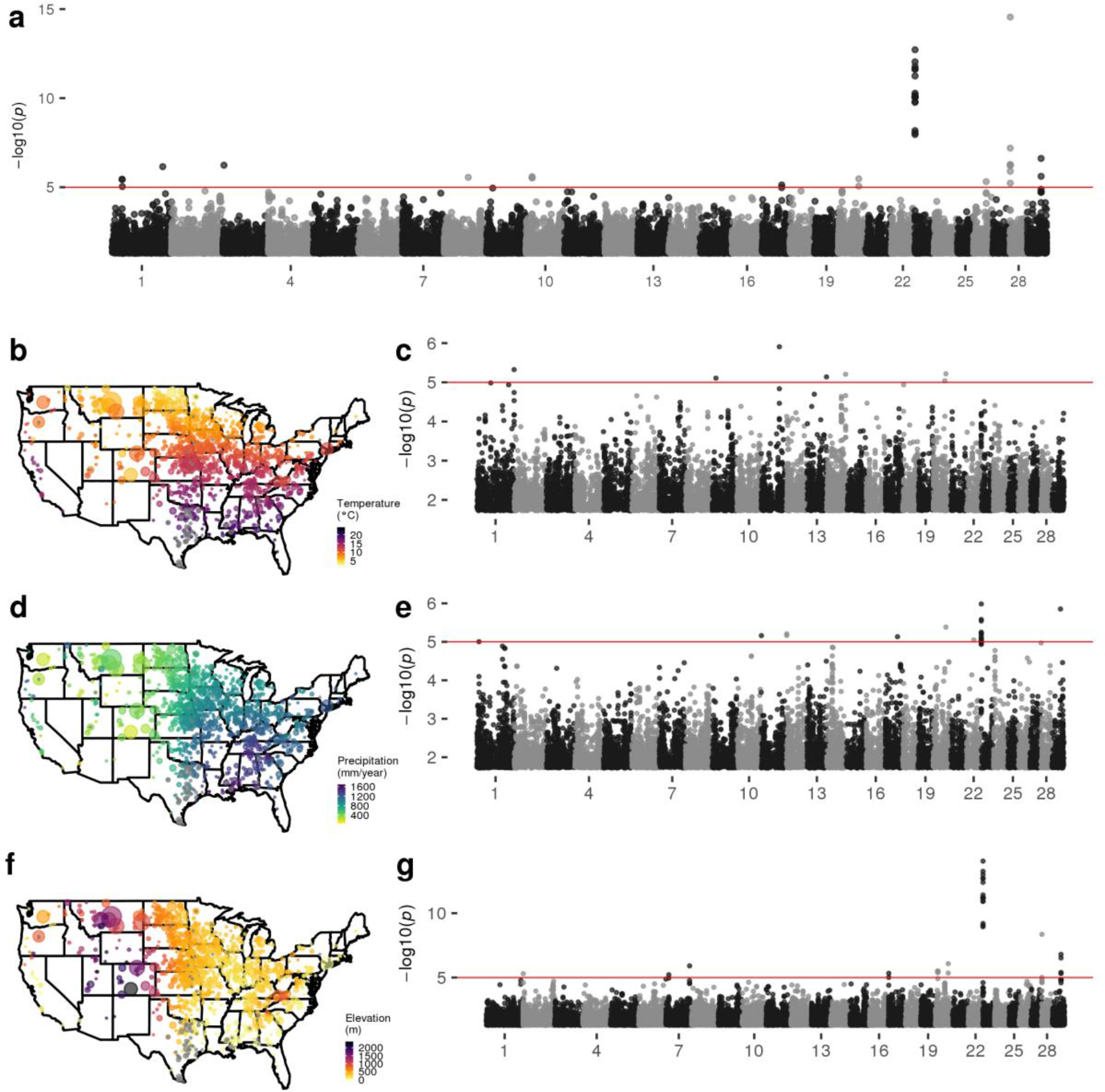
Continuous environmental variable envGWAS in Simmental cattle. (a) Multivariate envGWAS of temperature, precipitation, and elevation for Simmental cattle. Geographic distributions colored by (b) temperature, (d) precipitation, (f) elevation. Manhattan plots for univariate envGWAS analysis of (c) temperature, (e) precipitation, (g) elevation. Red lines indicate permutation-derived p-value cutoff of 1×10^-5^.

**Fig S9.**
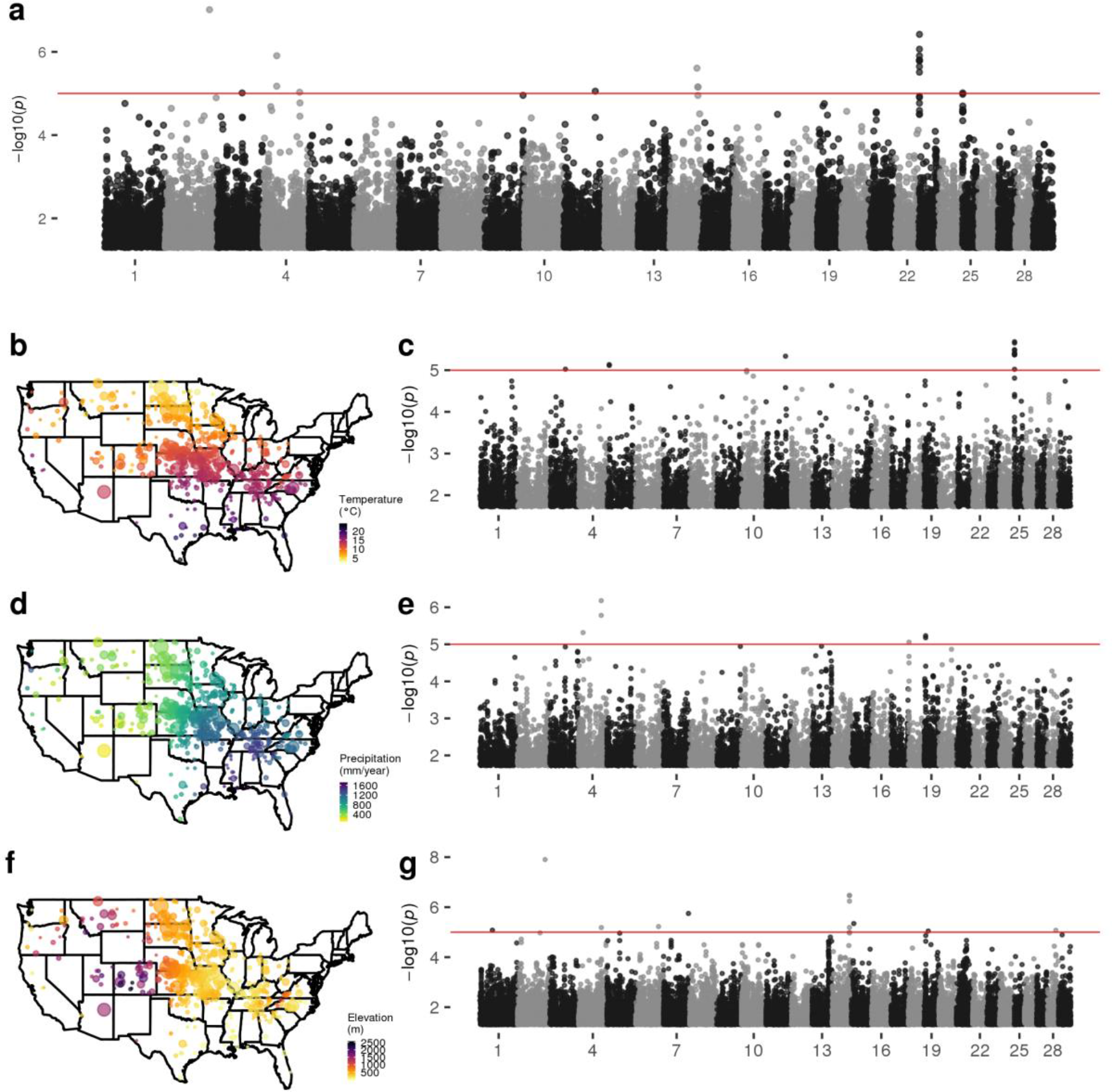
Continuous environmental variable envGWAS in Gelbvieh cattle. (a) Multivariate envGWAS of temperature, precipitation, and elevation for Gelbvieh cattle. Geographic distributions colored by (b) temperature, (d) precipitation, (f) elevation. Manhattan plots for univariate envGWAS analysis of (c) temperature, (e) precipitation, (g) elevation. Red lines indicate permutation-derived p-value cutoff of 1×10^-5^.

**Fig S10.**
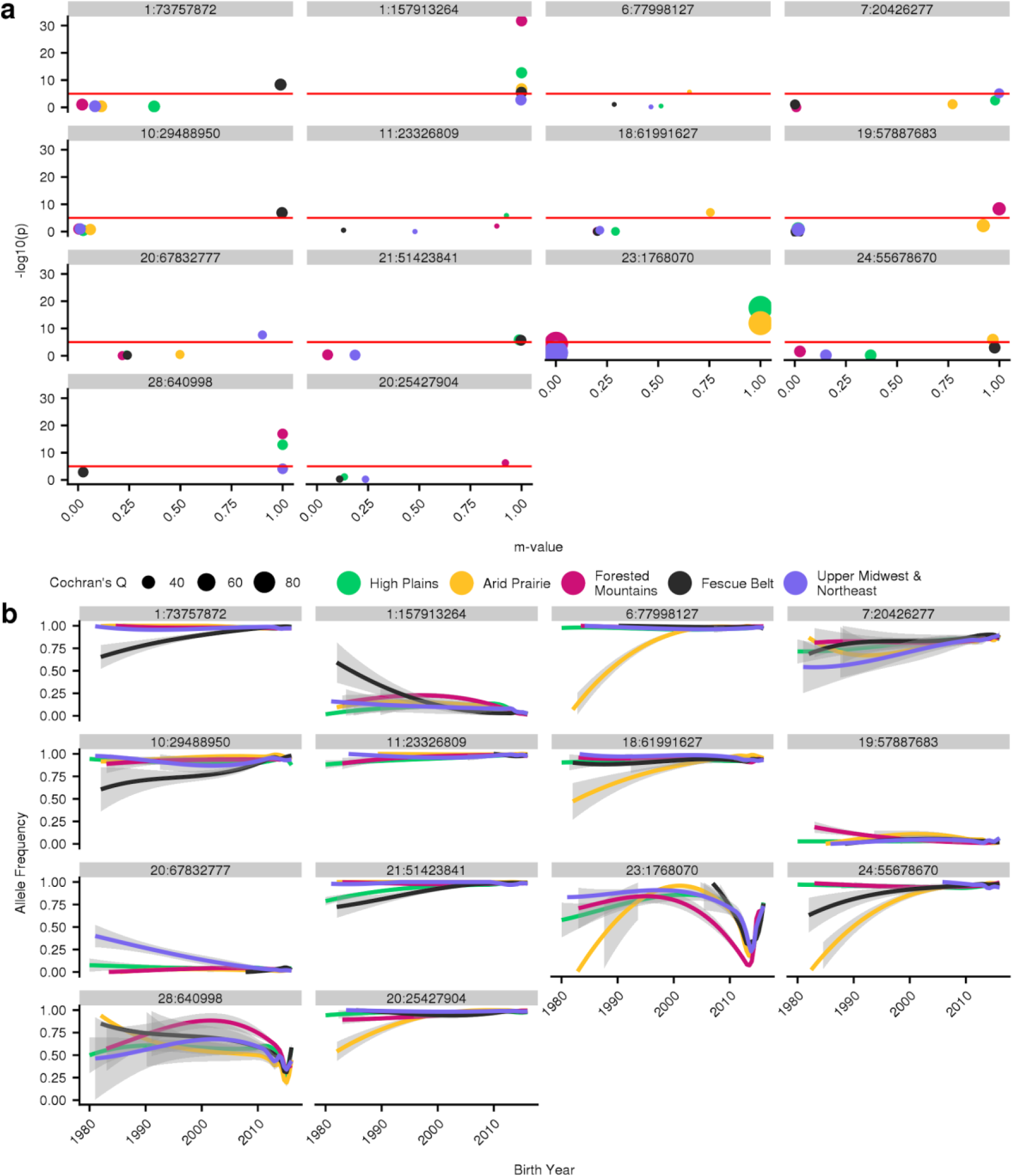
PM-plots and region-specific allele frequency trajectories for meta-analysis SNPs of interest in the Red Angus population ecoregions with > 1,000 genotyped animals. (a) PM-plots for lead SNPs of significant within-region GPSM meta-analysis (Cochran’s Q p-value > 1×10^-5^ and significant in at least one region-specific GPSM analysis p < 1×10^-5^). Each box represents the lead SNP, colored by ecoregion, and sized by Cochran’s Q value (for heterogeneity). (b) Region-specific allele frequency trajectories for lead SNPs since 1980, generated by fitting smoothed loess regression of allele frequency on birth date. Trajectories are colored by ecoregion.

**Fig S11.**
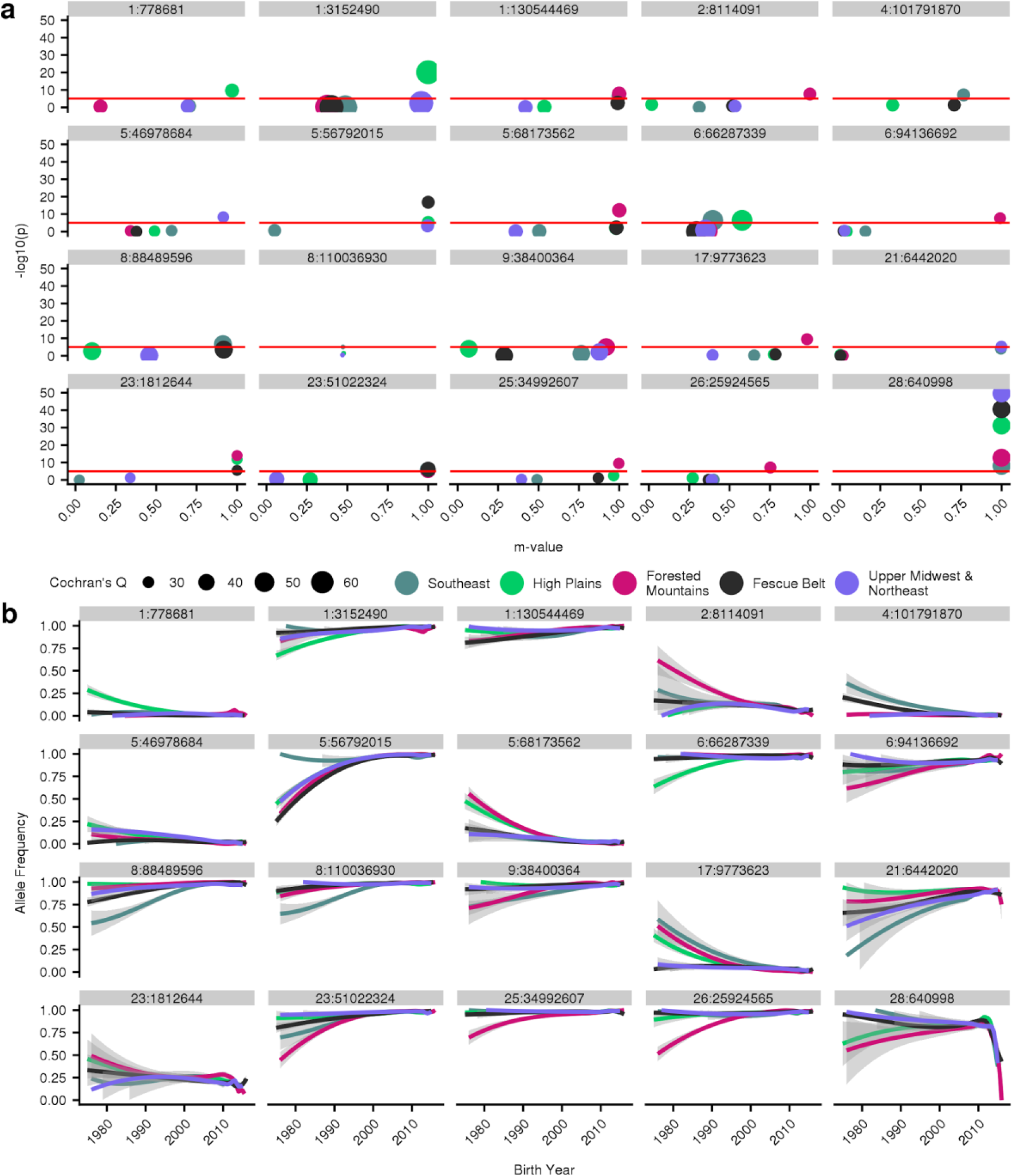
PM-plots and region-specific allele frequency trajectories for meta-analysis SNPs of interest in the Simmental population ecoregions with > 1,000 genotyped animals. (a) PM-plots for lead SNPs of significant within-region GPSM meta-analysis (Cochran’s Q p-value > 1×10^-5^ and significant in at least one region-specific GPSM analysis p < 1×10^-5^). Each box represents the lead SNP, colored by ecoregion, and sized by Cochran’s Q value (for heterogeneity). (b) Region-specific allele frequency trajectories for lead SNPs since 1980, generated by fitting smoothed loess regression of allele frequency on birth date. Trajectories are colored by ecoregion.

**Fig S12.**
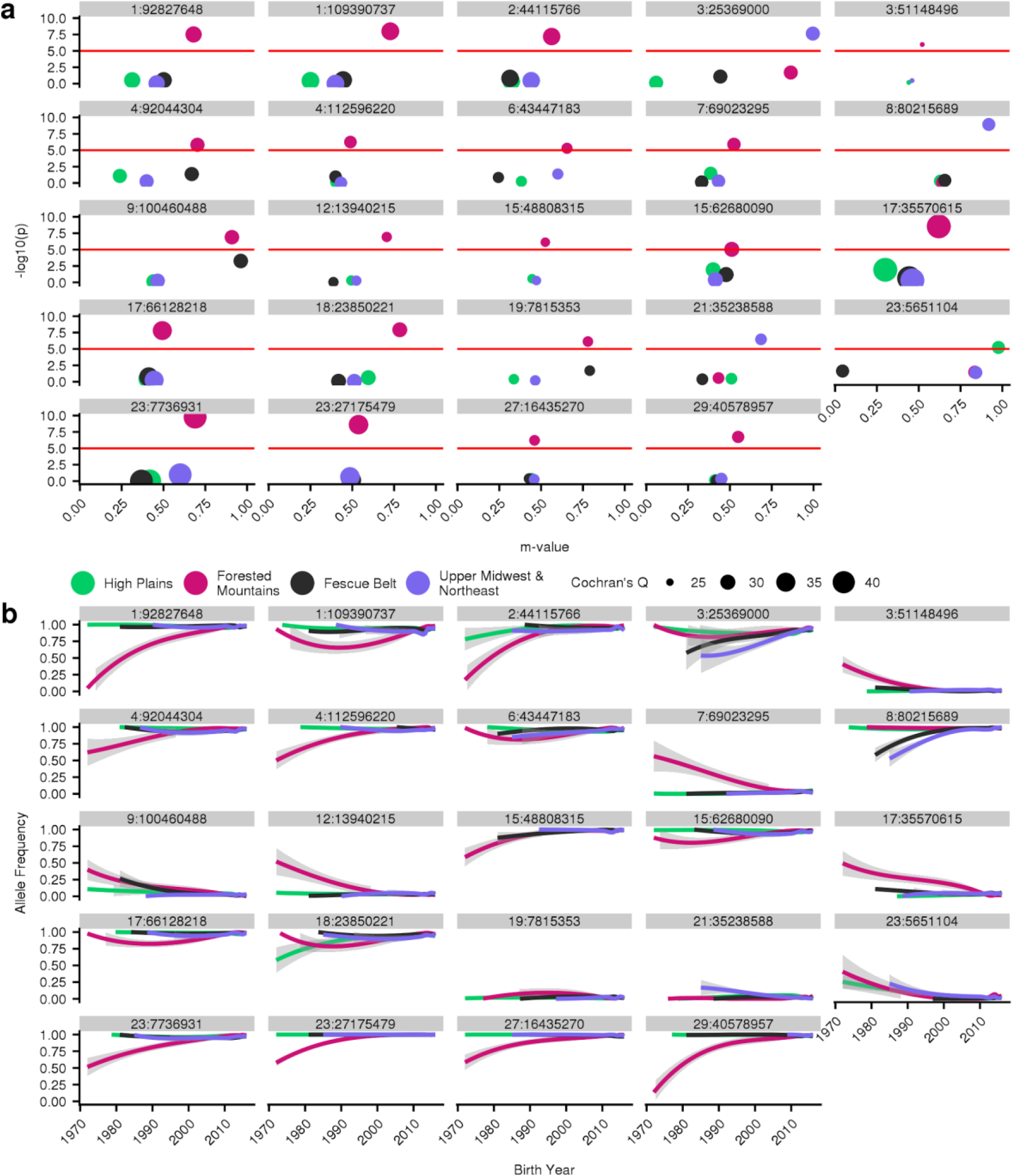
PM-plots and region-specific allele frequency trajectories for meta-analysis SNPs of interest in the Gelbvieh population ecoregions with > 1,000 genotyped animals. (a) PM-plots for lead SNPs of significant within-region GPSM meta-analysis (Cochran’s Q p-value > 1×10^-5^ and significant in at least one region-specific GPSM analysis p < 1×10^-5^). Each box represents the lead SNP, colored by ecoregion, and sized by Cochran’s Q value (for heterogeneity). (b) Region-specific allele frequency trajectories for lead SNPs since 1980, generated by fitting smoothed loess regression of birth of allele frequency on birth date. Trajectories are colored by ecoregion.

**Table S1.** GPSM and envGWAS gene dropping simulation results. Ten repetitions of a 200,000 SNP gene dropping experiment through the complete Red Angus pedigree. Real GPSM and envGWAS phenotypes were used to identify significant SNPs (q-value < 0.1). Multiple nearby SNPs were grouped as “genomic regions’’ when they were within 1Mb of one another.

| Analysis | Analyzed Individuals | Median Number Significant Loci (SD) | Median Significant q-value (SD) |
| --- | --- | --- | --- |
| GPSM | 15,315 | 0.4 (0.52) | 0.047 (0.027) |
| envGWAS | 15,315 | 1.36 (1.56) | 0.051 (0.031) |

**Table S2. GPSM stochastic simulation results.** Descriptions of 36 selection scenarios, and the corresponding true and false positive rates for GPSM detecting simulated QTL under selection (simulated QTL GPSM q-value < 0.1). Each scenario’s true and false positive statistics were calculated based on 10 replicates starting with different founder populations and selected randomly (false positives) or based on true breeding value (true positives). For each scenario, we report the number simulated QTL, the number of total crosses performed using 50 males and 500 females, the distribution from which QTL effects were drawn from, and the number of generations of selection performed. The mean PVE was reported for both selection on true breeding value (TBV) and random mating. We report results when 10,000 simulated individuals were randomly chosen to be genotyped (evenly each generation) and when more recent animals were genotyped more frequently (uneven sampling).

**Table S3.**
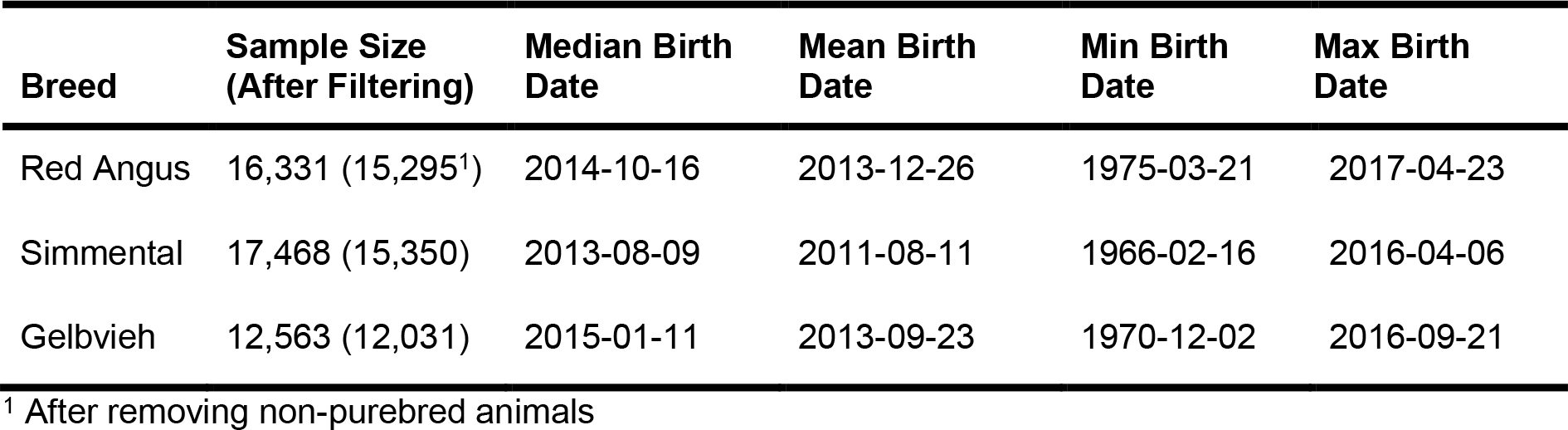
GPSM datasets from three major U.S. beef cattle populations. Sample sizes are reported prior to and after filtering on individual call rate individuals with reported birth dates.

| Breed | Sample Size (After Filtering) | Median Birth Date | Mean Birth Date | Min Birth Date | Max Birth Date |
| --- | --- | --- | --- | --- | --- |
| Red Angus | 16,331 (15,295<sup>1</sup>) | 2014-10-16 | 2013-12-26 | 1975-03-21 | 2017-04-23 |
| Simmental | 17,468 (15,350) | 2013-08-09 | 2011-08-11 | 1966-02-16 | 2016-04-06 |
| Gelbvieh | 12,563 (12,031) | 2015-01-11 | 2013-09-23 | 1970-12-02 | 2016-09-21 |

<sup>1</sup> After removing non-purebred animals

**Table S4.** The proportion of variation in birth date explained (PVE) by markers in GPSM analysis. PVE calculated for each population dataset in full, and subsetted to individuals born within the last 20 or 10 years. The standard errors of PVE estimates are reported in parentheses.

| Population | Full Dataset PVE (se) | 20-year PVE (se) | 10-year PVE (se) | Shuffled PVE (se) |
| --- | --- | --- | --- | --- |
| Red Angus | 0.520 (0.013) | 0.358 (0.013) | 0.406 (0.014) | $9.82 \times 10^{-6}$ (0.002) |
| Simmental | 0.588 (0.009) | 0.551 (0.010) | 0.401 (0.014) | $9.70 \times 10^{-4}$ (0.003) |
| Gelbvieh | 0.459 (0.015) | 0.454 (0.015) | 0.361 (0.014) | $5.05 \times 10^{-4}$ (0.004) |

**Table S5. Summary statistics and candidate genes from significant GPSM SNPs for Red Angus, Simmental, and Gelbvieh populations.** Variants are significant if GPSM q-value < 0.1. Genomic locations are reported based on coordinates from ARS-1.2 genome assembly. Candidate genes were assigned to a SNP if within 10 kb of a significant SNP.

**Table S6.** Summary statistics of allele frequency change (ΔAF) per generation for significant GPSM SNPs. ΔAF is the slope of a simple regression of allele frequency on birth date multiplied by a generation interval of 5 years.

| Breed | N SNPs (GPSM q < 0.1) | Mean ΔAF per generation (sd) | Median ΔAF per generation | Min ΔAF per generation | Max ΔAF per generation |
| --- | --- | --- | --- | --- | --- |
| Red Angus | 268 | 0.018 (0.011) | 0.017 | $4.97 \times 10^{-5}$ | 0.076 |
| Simmental | 548 | 0.024 (0.017) | 0.022 | $6.79 \times 10^{-5}$ | 0.093 |
| Gelbvieh | 762 | 0.033 (0.028) | 0.024 | $1.01 \times 10^{-4}$ | 0.223 |

**Table S7.**
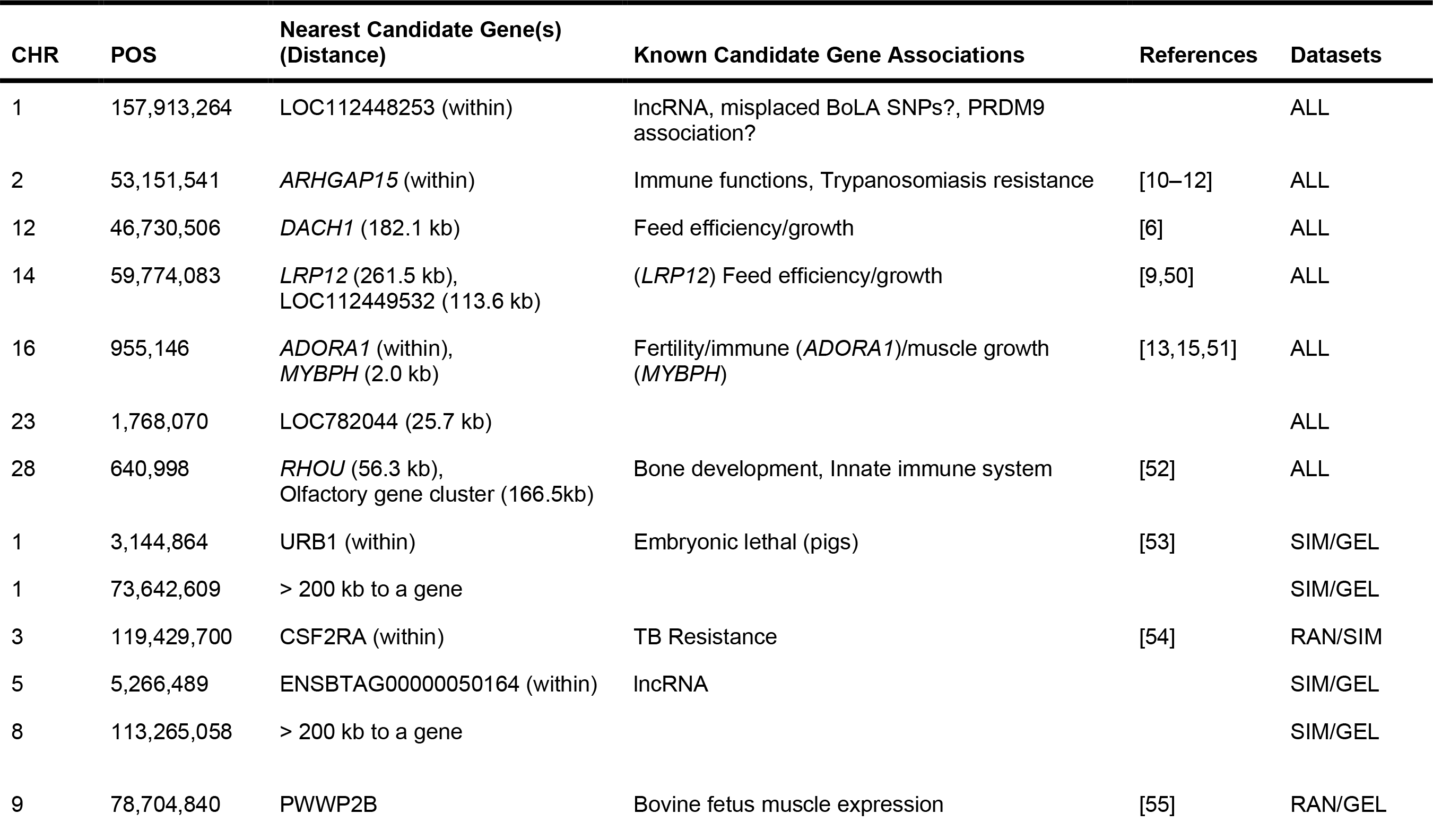
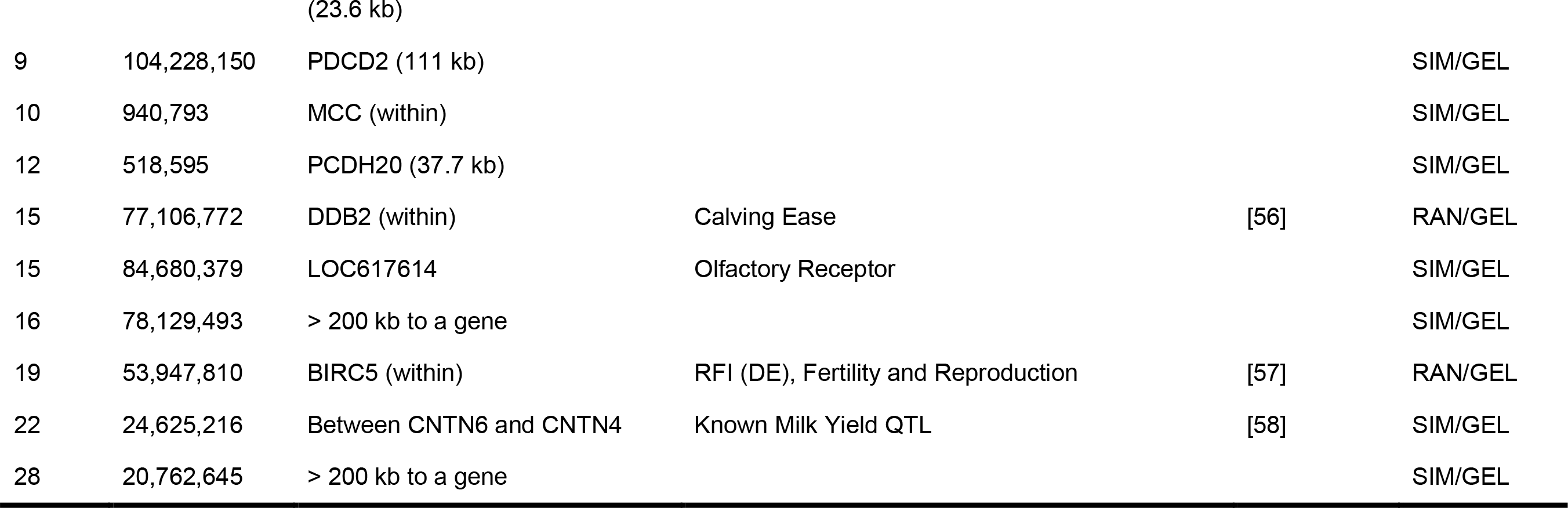
Significant GPSM variants identified in at least two populations. Lead SNPs from significant loci identified in GPSM analyses of Red Angus, Simmental, and Gelbvieh cattle populations. Locus reported if it was identified in GPSM analysis of at least two populations. Candidate gene is the annotated gene closest to lowest q-value SNP in peak if < 200 kb away. Associations are from cattle literature unless otherwise reported.

**Table S8. Gene enrichment analysis of GPSM candidate genes in Red Angus, Simmental, and Gelbvieh populations.** Candidate genes were annotated genes < 10 kb to significant GPSM SNPs (q < 0.1). Significant (FDR-corrected p-values < 0.1) KEGG pathways and GO biological processes are reported for each breed.

**Table S9. TissueEnrich analysis using GPSM gene sets from Red Angus, Simmental, and Gelbvieh populations.** TSEA from TissueEnrich software using Human Protein Atlas gene expression data. Enrichment analysis carried out for candidate genes within 10 kb of significant envGWAS SNPs. For each test, we report the number of tissue specific genes, their average fold change, and the FDR-corrected log_10_ p-value for tissue enriched expression.

**Table S10. Tissue enrichment analysis results from GPSM gene sets in Red Angus, Simmental, and Gelbvieh populations using the pSI R package and human GTEx expression data.** Enrichment significance values for four specificity index thresholds (pSI) of 25 human tissues types. Each combination of stringency for enrichment (pSI) and tissue reports a p-value for Fisher’s Exact Test and a Benjamini Hochberg corrected p-value reported in parentheses. Tissue-gene-set combinations that are significant (Benjamini-Hochberg p-value < 0.1) are highlighted in red, those that are suggestive (raw p-value < 0.1) are highlighted in green.

**Table S11.**
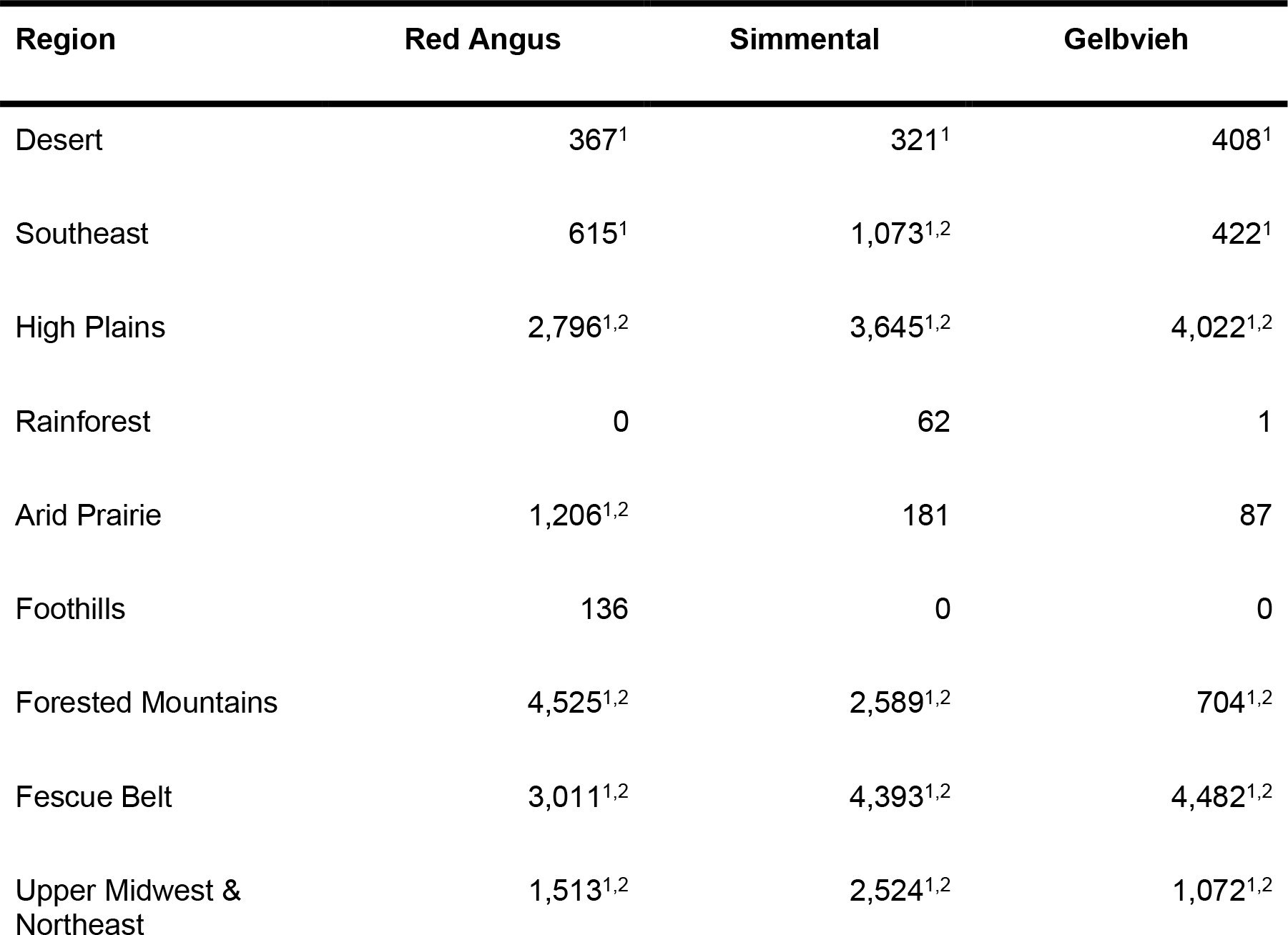
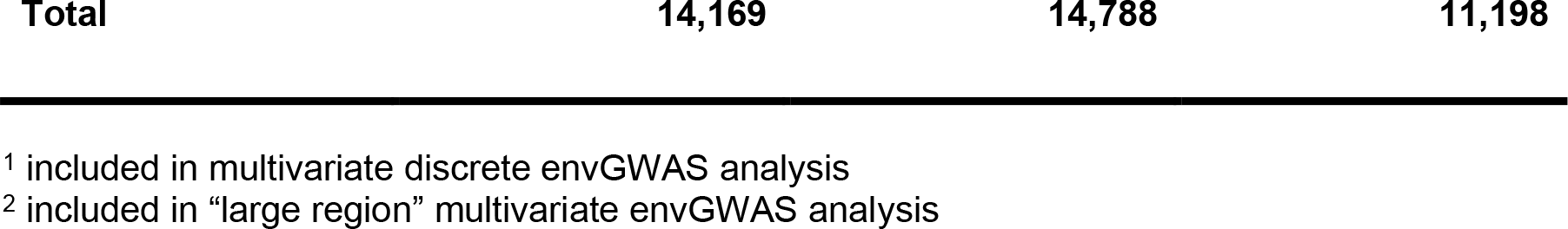
Ecoregion distribution of Red Angus, Simmental, and Gelbvieh populations. Counts of analyzed individuals in each region for each dataset after filtering and region assignment based on individual’s breeder zip code.

| Region | Red Angus | Simmental | Gelbvieh |
| --- | --- | --- | --- |
| Desert | 367<sup>1</sup> | 321<sup>1</sup> | 408<sup>1</sup> |
| Southeast | 615<sup>1</sup> | 1,073<sup>1,2</sup> | 422<sup>1</sup> |
| High Plains | 2,796<sup>1,2</sup> | 3,645<sup>1,2</sup> | 4,022<sup>1,2</sup> |
| Rainforest | 0 | 62 | 1 |
| Arid Prairie | 1,206<sup>1,2</sup> | 181 | 87 |
| Foothills | 136 | 0 | 0 |
| Forested Mountains | 4,525<sup>1,2</sup> | 2,589<sup>1,2</sup> | 704<sup>1,2</sup> |
| Fescue Belt | 3,011<sup>1,2</sup> | 4,393<sup>1,2</sup> | 4,482<sup>1,2</sup> |
| Upper Midwest & Northeast | 1,513<sup>1,2</sup> | 2,524<sup>1,2</sup> | 1,072<sup>1,2</sup> |
| **Total** | **14,169** | **14,788** | **11,198** |

<sup>1</sup> included in multivariate discrete envGWAS analysis
<sup>2</sup> included in “large region” multivariate envGWAS analysis

**Table S12.** Univariate REML estimates of PVE for continuous environmental variables in genotyped Red Angus, Simmental, and Gelbvieh populations. Standard errors for PVE estimates are reported in parentheses.

| Variable | Red Angus PVE (se) | Simmental PVE (se) | Gelbvieh PVE (se) |
| --- | --- | --- | --- |
| Temperature | 0.597 (0.010) | 0.586 (0.011) | 0.691 (0.010) |
| Precipitation | 0.526 (0.011) | 0.602 (0.011) | 0.677 (0.010) |
| Elevation | 0.594 (0.010) | 0.585 (0.011) | 0.644 (0.011) |

**Table S13.**
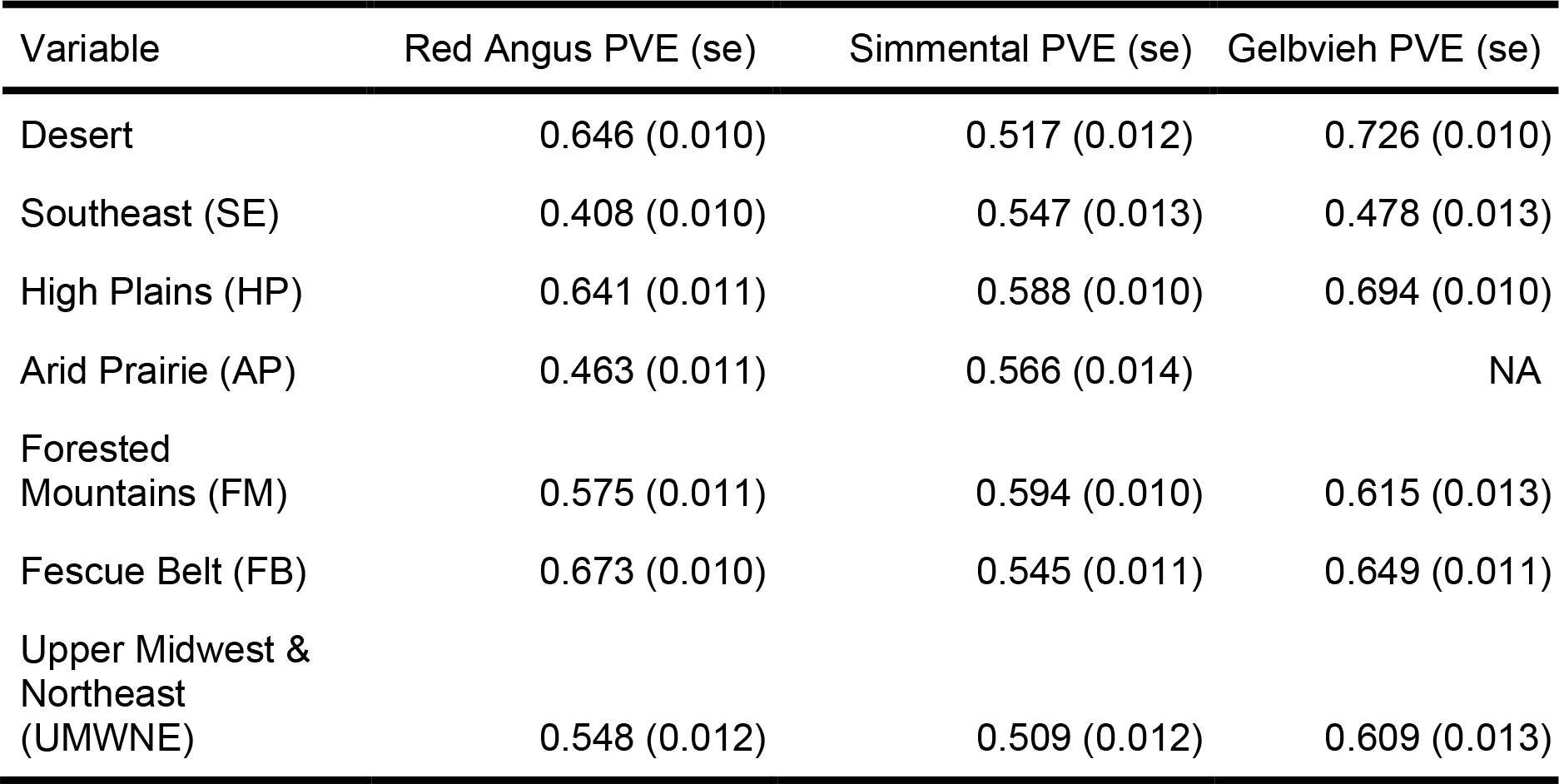
Univariate estimates of PVE for discrete ecoregion assignment in genotyped Red Angus, Simmental, and Gelbvieh populations. Standard errors for PVE estimates are reported in parentheses.

**Table S14.** Candidate genes for discrete ecoregion multivariate envGWAS in Red Angus cattle. Lead SNP in envGWAS peak is reported along with nearest plausible candidate genes (provided < 250 kb from lead SNP). If association was also identified in univariate analysis, it is reported. Potentially adaptive associations are reported along with references.

| CHR | POS | Nearest Candidate Gene(s) (Distance) | Univariate Continuous Association | Univariate Ecoregion Association | Candidate Gene Adaptive Associations | Reference | Breed(s) |
| --- | --- | --- | --- | --- | --- | --- | --- |
| 2 | 7,571,508 | DIRC1 (110.3 kb), COL5A2 (212.7 kb) | Multivariate, Temperature | FM | Sweep region (human), blood pressure (human) | [25,59,60] | RAN |
| 2 | 25,519,381 | GORASP2 (4.47 kb) | N/A | AP | Adaptive signature (fish) | [61] | RAN |
| 4 | 43,928,337 | MAGI2 (within) | N/A | UMWNE | Selection signature, imprinted (cattle) | [62,63] | RAN |
| 5 | 59,498,938 | LOC788524 (OR9K2) (1.42 kb) [Olfactory cluster] | N/A | FM | Olfactory receptor cluster, selection signature (bison) | [20] | RAN |
| 6 | 96,217,506 | RASGEF1B (within) | N/A | HP | Immune function, adaptation signature (human), Response to viral infections. | [64,65] | RAN |
| 8 | 84,146,584 | CENPP (within) | N/A | N/A | African cattle CNV, hypoxia (human) | [66,67] | RAN |
| 12 | 58,438,020 | N/A | N/A | N/A |  |  | RAN |
| 13 | 75,784,164 | ZMYND8 (within) |  |  | Immune function, DNA damage repair | [68,69] | RAN |
| 13 | 81,716,869 | PFDN4 (71.97 kb) | N/A | HP | Dermatitis (humans) | [70] | RAN |
| 22 | 48,873,180 | DUSP7 (40.83 kb) |  |  | Heat stress/sperm motility (cattle), Stress response | [71,72] | RAN |
| 23 | 1,768,070 |  |  | AP, FM |  |  | RAN |
| 24 | 10,628,280 | CDH7 (151.7 kb) | MV | AP | Developmental processes | [73] | RAN |
| 24 | 62,228,300 | LOC100298064<br>(HAVCR1) (15.2kb) |  | AP | Immune function | [74] | RAN |
| 25 | 26,511,450 | SPN (CD43) (within) | N/A | SE | MHC-Class I, Immune<br>response (cattle) | [75] | RAN |
| 25 | 35,085,041 | CUX1 (67.8 kb) | Multivariate | N/A | Hair phenotype (goats, mice) | [76,77] |  |
| 26 | 34,432,722 | ADRB1 (81.5 kb) | Temperature |  | Selection (sporting dogs),<br>climate adaptation (lizards) | [78,79] | RAN |

**Table S15. envGWAS significant SNPs and candidate genes.** SNPs were significant when p < 1×10^-5^. Candidate genes are genes within < 10 kb of significant envGWAS SNPs. We report significant SNPs from all univariate and multivariate analyses for both continuous environmental variables and discrete environments in Red Angus, Simmental, and Gelbvieh populations.

**Table S16.**
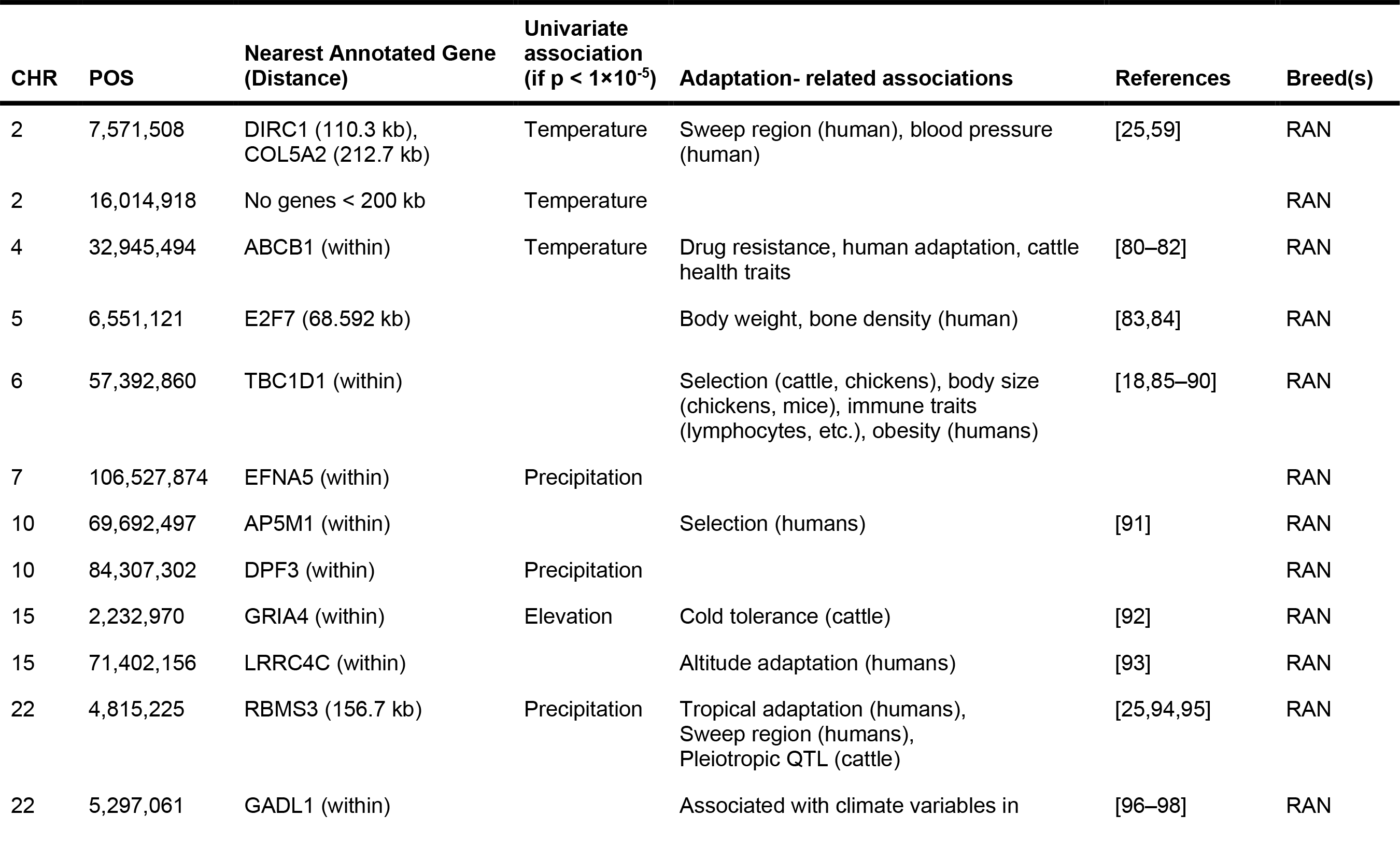
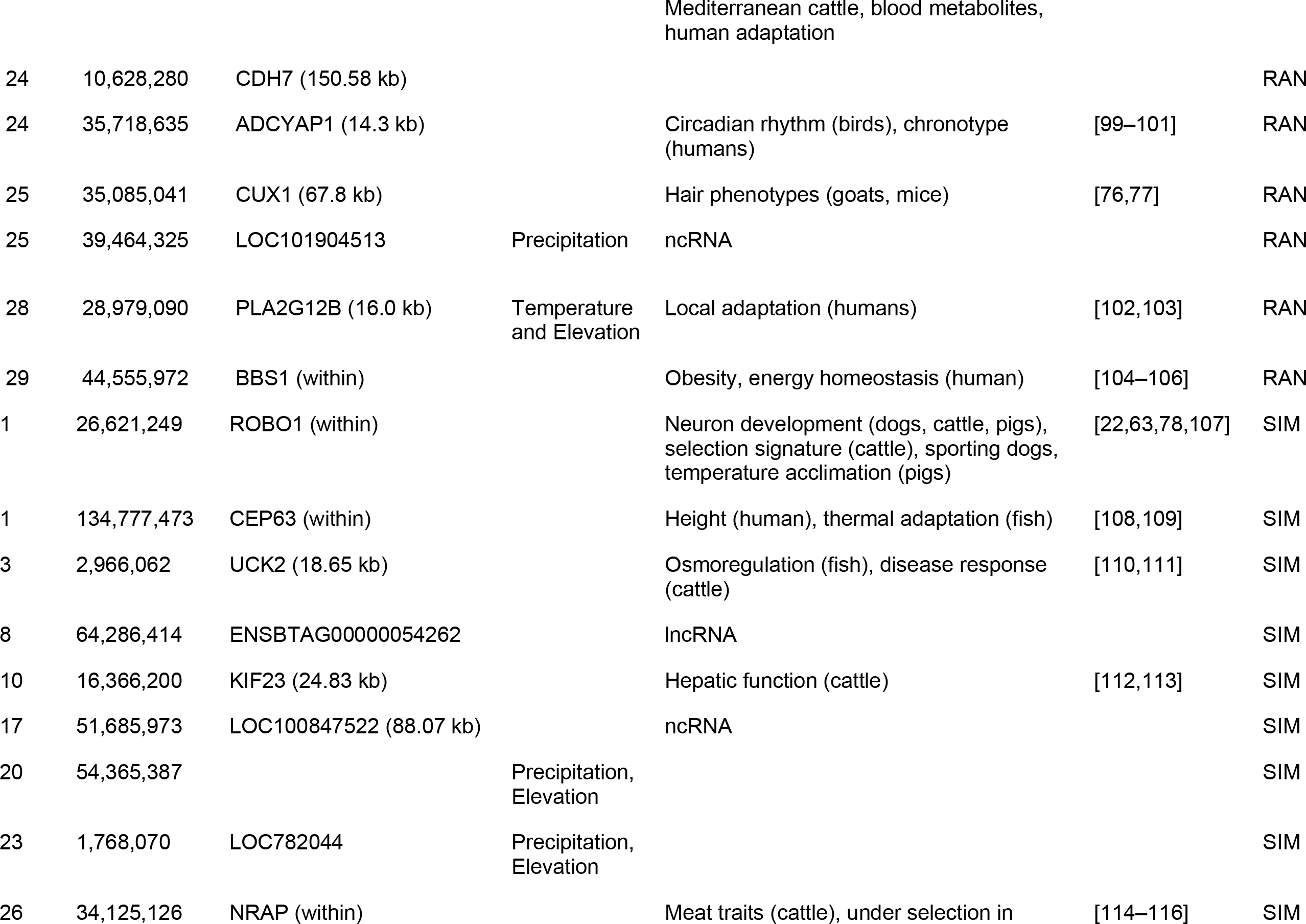
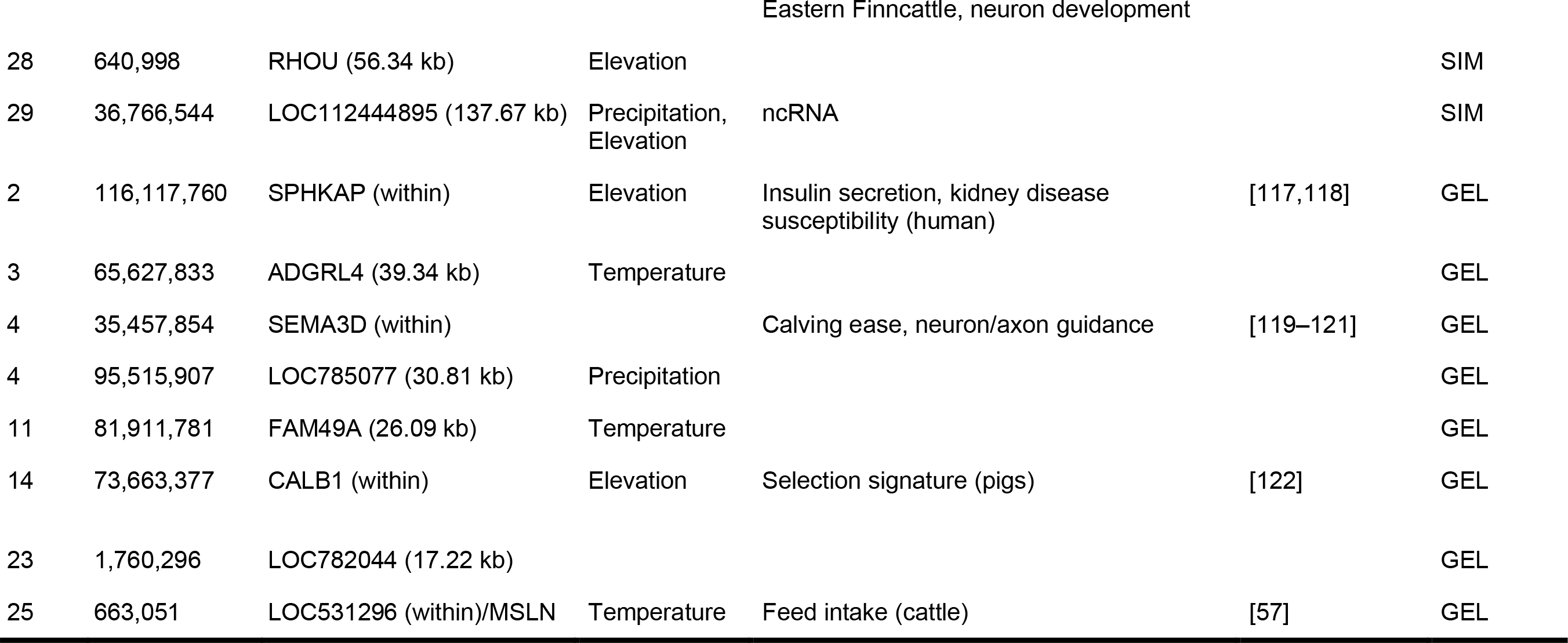
Candidate genes identified in multivariate envGWAS analyses using continuous environmental attributes as dependent variables. Chromosome and genomic positions are for lead SNP in peak. Closest gene is identified as a candidate (if < 250 kb from lead SNP).

**Table S17. Gene enrichment analysis of envGWAS candidate genes in Red Angus, Simmental, and Gelbvieh populations.** Candidate genes were annotated genes < 10 kb to significant envGWAS SNPs (p-value < 1×10^-5^). A single gene list for each breed was generated using significant SNPs from all combinations of univariate/multivariate, continuous/discrete envGWAS. Significant (FDR-corrected p-values < 0.1) KEGG pathways and GO biological processes are reported, along with the associated genes.

**Table S18. TissueEnrich analysis using envGWAS gene sets from Red Angus, Simmental, and Gelbvieh populations.** TSEA from TissueEnrich software using Human Protein Atlas gene expression data. Enrichment analysis carried out for candidate genes within 10 kb of significant envGWAS SNPs. For each test, we report the number of tissue specific genes, their average fold change, and the FDR-corrected log_10_ p-value for tissue enriched expression.

**Table S19. Tissue enrichment analysis results from envGWAS gene sets in Red Angus, Simmental, and Gelbvieh populations using the pSI R package and human GTEx expression data.** Enrichment significance values for four specificity index thresholds (pSI) of 25 human tissues types. Each combination of stringency for enrichment (pSI) and tissue reports a p-value for Fisher’s Exact Test and a Benjamini Hochberg corrected p-value reported in parentheses. Tissue-gene-set combinations that are significant (Benjamini-Hochberg p-value < 0.1) are highlighted in red, those that are suggestive (raw p-value < 0.1) are highlighted in green.

**Table S20. Brain region and cell-type enrichment analysis results from envGWAS gene sets in Red Angus, Simmental, and Gelbvieh populations using the pSI R package with expression data from the Allen Brain Atlas.** Enrichment significance values for four specificity index thresholds (pSI) of six brain regions and 35 brain cell types. Each combination of stringency for enrichment (pSI) and brain region/cell type reports a p-value for Fisher’s Exact Test and a Benjamini Hochberg corrected p-value reported in parentheses. Combinations that are significant (Benjamini-Hochberg p-value < 0.1) are highlighted in red, those that are suggestive (raw p-value < 0.1) are highlighted in green.

**Table S21. Cell-type specific expression of homologous *C. elegans* genes derived from envGWAS candidate gene lists of Red Angus, Simmental, and Gelbvieh populations.** *C. elegans* gene homologs were generated from Ortholist2, requiring that genes be present in at least three data sources to be included in enrichment analysis. For each breed’s gene list, we include a list of worm tissues with significant enrichment of listed genes (q-value < 0.1).

